# Optoelectronic enhancement of photocurrent by cyanobacteria on sustainable AP-VPP-fabricated PEDOT electrodes

**DOI:** 10.1101/2023.10.11.561827

**Authors:** Laura T. Wey, Rahul Yewale, Emilia Hautala, Jenna Hannonen, Kalle Katavisto, Carita Kvarnström, Yagut Allahverdiyeva, Pia Damlin

**Affiliations:** Photomicrobes Research Group, Molecular Plant Biology unit, Department of Life Technologies, University of Turku, 20014 Turku, Finland; Materials Chemistry Research Group, Department of Chemistry, University of Turku, 20014 Turku, Finland

**Author notes:** Laboratory of Industrial Physics, Department of Physics and Astronomy, University of Turku, 20014 Turku, Finland (Rahul Yewale). Battery Materials and Technologies Research Group, Department of Mechanical and Materials Engineering, University of Turku, FI-20014 Turun yliopisto, Finland.

**Keywords:** Biophotovoltaics, conducting polymers, vapor phase polymerization, photosynthetic microorganisms, exoelectrogenesis

## Abstract

Photosynthetic microrganisms, including cyanobacteria, can be interfaced with electrodes in biophotovoltaic devices (BPVs) for solar energy conversion. Effective BPV electrodes need to be conductive, transparent, flexible, biocompatible and environmentally friendly, while also being cost-effective, abundant in material and lightweight. The utilization of electrically conducting polymers (CPs), particularly poly(3,4-ethylenedioxythiophene) (PEDOT) fabricated by an atmospheric pressure vapor phase polymerization (AP-VPP) technique, is a promising avenue for BPV applications. However, challenges remain in optimising their performance as CPs are dynamic optoelectronic materials, and their interaction with photosynthetic biocatalysts under a range of conditions has not been explored thoroughly. Here we show that AP-VPP-PEDOT electrodes hold promise for interfacing with cyanobacteria in BPVs to generate green electricity under red and blue light and moderate applied potentials with exogenous electron mediators. The highest non-mediated photocurrent achieved was 0.48 µA cm^−2^, with a two-layer PEDOT electrode at 0.5 V applied potential and blue light. The highest mediated photocurrent achieved was 2.73 µA cm^−2^, with a one-layer PEDOT electrode at 0.3 V applied potential and blue light and the exogenous electron mediator 2,6-dichloro-1,4-benzoquinone (DCBQ). The proposed approach to fabricating PEDOT electrodes offers a new pathway for developing sustainable electrodes for BPVs and pinpoints strategies for future optimisation for achieving high-performance outcomes.

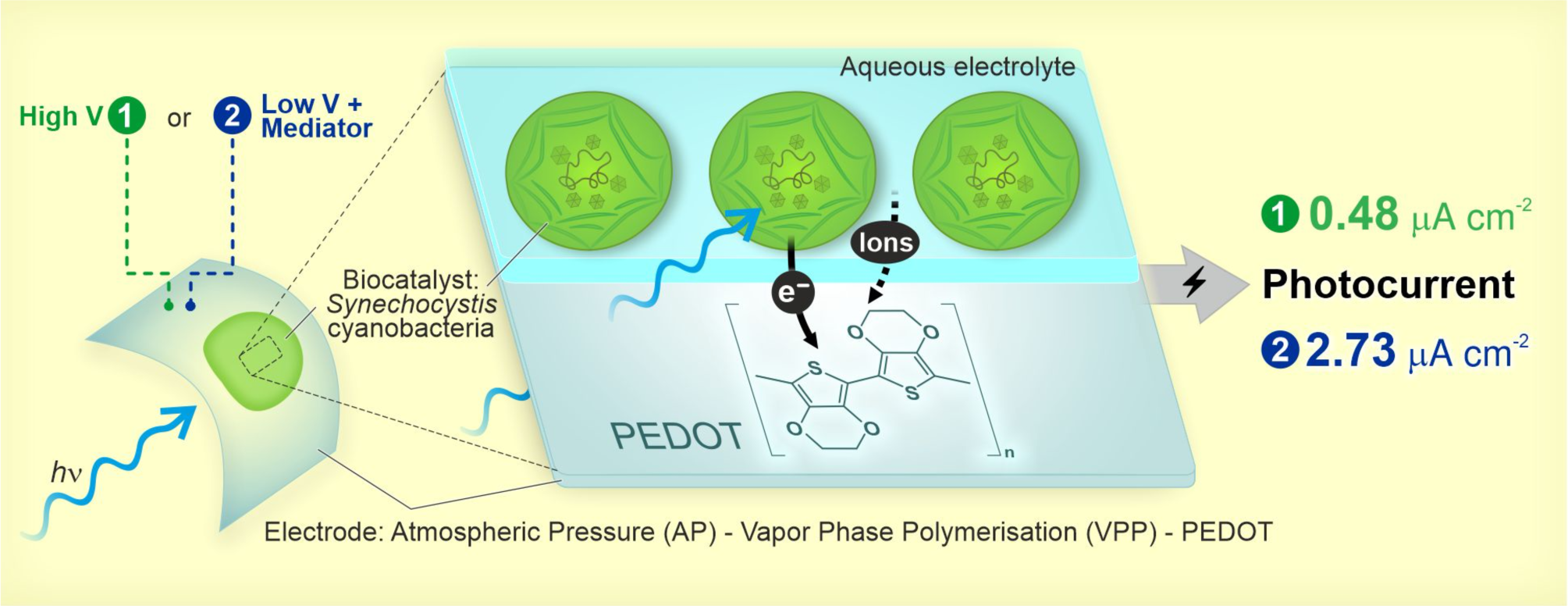

## 1. Introduction

In light of the severe energy crisis caused by climate change and the depletion of natural resources, the need for clean, renewable energy sources has become more urgent than ever.[1] Biophotovoltaic devices (BPVs) are an emerging biotechnology that generate electricity from photosynthetic microorganisms (algae and cyanobacteria) performing exoelectrogenesis under illumination as a by-product of photosynthesis (‘photocurrent’, **Figure 1**).[2,3] BPVs have been modelled to be able to power devices in remote locations, the Internet of Things and even be integrated onto buildings; recently, they have powered a microprocessor for at least six months.[4–6] However, the electrodes used to achieve present benchmark photocurrent outputs of 1.93 µA cm^−2^ (245 µA cm^−2^ with an exogenous electron mediator)[7] are made of indium tin oxide (ITO), where indium is a rare earth metal[8,9] and ITO nanoparticles have toxicity-associated bioaccumulation.[10–12] To achieve truly green electricity generation, environmentally friendly and non-toxic electrodes are needed.

**Figure 1.**
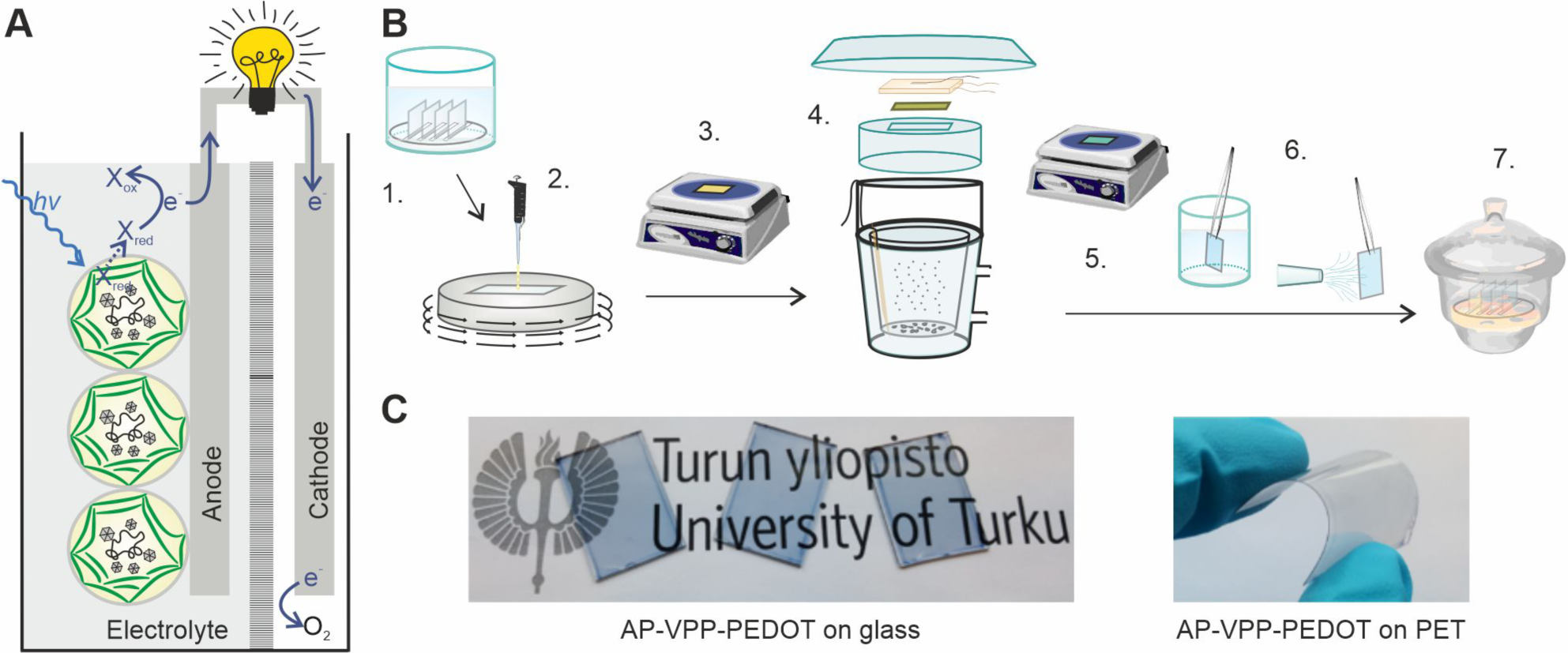
Schematic overview. **A)** A biophotovoltaic device (BPV) for power generation from exoelectrogenesis of photosynthetic microorganisms interfaced with an anode electrode under illumination. X is an endogenous electron mediator exported by the cell in its reduced form (Xred) and oxidized at the electrode surface (Xox) generating current.[63,64] **B)** Various steps in fabricating PEDOT thin films using optimized vapor phase polymerization at atmospheric pressure (AP-VPP):[16,58,21,59] 1. Substrate cleaning, 2. Spin coating of oxidant solution, 3. Oxidant film drying, 4. Polymerization, 5. Annealing, 6. Washing the films, 7. Storage prior to application. **C)** Photos of AP-VPP PEDOT film on glass showing transparency and on polyethylene terephthalate (PET) showing flexibility.

The BPV community can look to the commercial flexible display screen/touch screen technology field for inspiration, as they are shifting away from ITO electrodes and there is alignment in their desirable electrode material properties: conductive, transparent, flexible, biocompatible and environmentally friendly, as well as low cost, material abundance and lightweight. Electrically conducting polymers (CPs, also called conjugated polymers, synthetic metals or organic semiconductors), are used as electrodes in the screen field [14] and are promising material candidates for BPVs. CPs are polymers containing conjugated sp^2^ carbons with low-tuneable bandgap and overlapping molecular orbitals with delocalized π-electrons that allow transport of charge.[15]

Poly(3,4-ethylenedioxythiophene) (PEDOT) is a promising CP for BPV applications (**Table 1**). PEDOT has been extensively studied due to its high, stable, and tuneable electrical conductivity (up to 4600 ± 100 S cm^-1^)[16–21] that can be varied with the type of dopant ion and polymerization method. PEDOT is transparent if made into thin films,[22] which makes it ideal for light management for photosynthetic biocatalysts. Light intensity throughout the bio-anode has recently been identified as the bottleneck on the non-biological-side of optimising BPVs.[7] PEDOT can be made on flexible substrates and does not lose its conductivity upon bending,[23] making it ideal for upscaling, construction and even wearable technology.[24] CPs like PEDOT are environmentally friendly and biodegradable, some CP monomers such as azulene naturally occur in plants and mushrooms.[25] While toxicity studies of CP monomers are not intensively reported, some studies report EDOT monomer as less toxic than ITO nanoparticles[26,27] and PEDOT has been used in medical applications.[28] For BPV applications it is imperative to additionally ensure a material’s biocompatibility with photosynthetic cells; a PEDOT–PSS composite nanofiber with cellulose nanofibrils and poly(ethylene oxide) has been shown to not inhibit the photosynthetic efficiency of the photosynthetic pigment-protein complex, photosystem II (PSII), in the model green algae and cyanobacteria cells.[29]

**Table 1.**
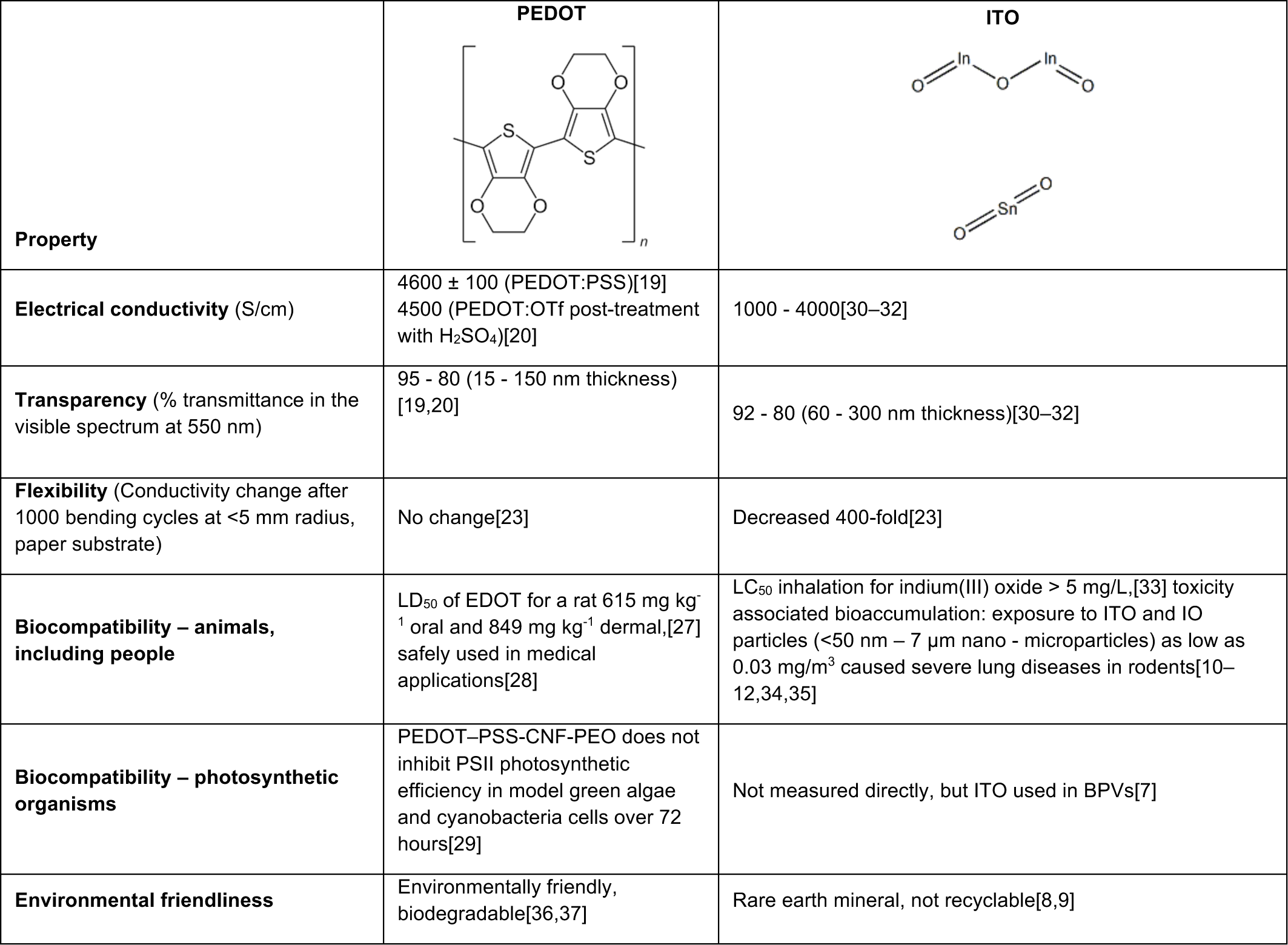
Properties of candidate conducting polymer poly(3,4-ethylenedioxythiophene) (PEDOT) compared to commonly used indium tin oxide (ITO). Abbreviations: Poly(styrene sulfonate) (PSS), trifluoromethanesulfonate (OTf), cellulose nanofibrils (CNF) and poly(ethylene oxide) (PEO), photosystem II (PSII).

| Property | PEDOT | ITO |
| --- | --- | --- |
| **Electrical conductivity** (S/cm) | 4600 ± 100 (PEDOT:PSS)[19]<br>4500 (PEDOT:OTf post-treatment with H<sub>2</sub>SO<sub>4</sub>)[20] | 1000 - 4000[30–32] |
| **Transparency** (% transmittance in the visible spectrum at 550 nm) | 95 - 80 (15 - 150 nm thickness) [19,20] | 92 - 80 (60 - 300 nm thickness)[30–32] |
| **Flexibility** (Conductivity change after 1000 bending cycles at <5 mm radius, paper substrate) | No change[23] | Decreased 400-fold[23] |
| **Biocompatibility – animals, including people** | LD<sub>50</sub> of EDOT for a rat 615 mg kg<sup>-1</sup> oral and 849 mg kg<sup>-1</sup> dermal,[27] safely used in medical applications[28] | LC<sub>50</sub> inhalation for indium(III) oxide > 5 mg/L,[33] toxicity associated bioaccumulation: exposure to ITO and IO particles (<50 nm – 7 µm nano - microparticles) as low as 0.03 mg/m<sup>3</sup> caused severe lung diseases in rodents[10–12,34,35] |
| **Biocompatibility – photosynthetic organisms** | PEDOT–PSS–CNF–PEO does not inhibit PSII photosynthetic efficiency in model green algae and cyanobacteria cells over 72 hours[29] | Not measured directly, but ITO used in BPVs[7] |
| **Environmental friendliness** | Environmentally friendly, biodegradable[36,37] | Rare earth mineral, not recyclable[8,9] |

In previous studies, CPs have been used to coat the surface of an electrode substrate (such as ITO, carbon cloth or graphite) in microbial fuel cells (MFCs),[38–48] and, more recently, in BPVs (**Supplementary Table 1**).[49–56] In these cases, the CP acts as a mediator between the exoelectrogenic microorganisms and the electrode to boost the photocurrent and power outputs.[57] The CP also adds nanoroughness and positive charges to the surface of the electrode substrate to enhance the adhesion of negatively charged cells.[56] PEDOT can be manufactured in different forms using different methods. PEDOT:PSS polymer composite is the form of PEDOT that has been used in previous BPV studies, where it was brush painted onto carbon cloth[19] or fabricated by electrochemical polymerization on graphite rods using sodium dodecyl sulfate (SDS) as dopant ion yielding cloud-like thick polymer films.[55] In PEDOT:PSS, PSS, the stabilizer used to disperse PEDOT in the solvent, is an electrical insulator with high acidity. The intrinsically acidic nature of PEDOT:PSS causes deterioration and dewetting of the material, while its inhomogeneous conductivity and high surface roughness reduce the performance and stability of the electrodes.

We recently reported a new method to fabricate highly conducting PEDOT thin films onto non-conducting substrates (**Figure 1B**).[16,58,21,59] In brief, in the AP-VPP method an oxidant-coated substrate is exposed to monomer vapour at atmospheric pressure to form a thin layer of the polymer. After polymerization, a substrate containing a polymer film is annealed to avoid stress-fracture, and further washed and dried to get rid of excess amount of oxidant, unreacted monomer, reaction by-products and soluble species. Although PEDOT has previously been made by electrochemical synthesis, chemical synthesis, chemical vapor deposition, dip/spin/shear coating, and VPP under vacuum, these methods have limitations such as high electrode roughness, low conductivity, surface selectivity, high-temperature requirements, long polymerization time, and difficulties in achieving thin films, the latter of which is a key property to achieve transparency.[60–62] Moreover, since AP-VPP PEDOT can be fabricated on non-conducting substrate, the obtained photocurrent is purely due to cyanobacteria-CP interactions, enabling deconvolution of contribution from the substrate material.

In this paper we fabricated an electrode entirely made by the AP-VPP technique producing thin PEDOT films on nonconducting substrate to interface with cyanobacteria and harvest photocurrent for future applications in BPVs. We tested its electrochemical performance in an electrolyte compatible with the model cyanobacteria *Synechocystis* sp. PCC 6803 (hereafter *Synechocystis*). The biocompatibility of PEDOT films with *Synechocystis* was examined considering photosynthetic efficiency and growth. We systematically tested the photoelectrochemical performance of *Synechocystis* loaded on our AP-VPP-fabricated PEDOT electrodes under various key parameters, including cell loading, applied electrode potential, light wavelength, and PEDOT film layer thickness. Additionally, we measured the effect of addition of exogenous electron mediators on enhancing the efficiency of our device. We compared our electrodes to commercially available ITO electrodes and those reported in the literature. Furthermore, all measurements were performed with the light passing through the transparent AP-VPP-fabricated PEDOT electrodes.

Overall, our results demonstrate that AP-VPP-fabricated PEDOT holds promise as an anode material for interfacing with photosynthetic cyanobacteria *Synechocystis* in BPVs to generate green electricity. The proposed approach to fabricating PEDOT electrodes offers a new pathway for developing sustainable electrodes for BPVs. Furthermore, we have pinpointed strategies for future optimisation for achieving high-performance outcomes.

## 2. Experimental

### 2.1. Materials

#### 2.1.1. Chemicals

Iron(III) p-toluenesulfonate hexahydrate (FeTOS) (technical grade), tetrabutylammonium tetrafluoroborate (TBA-BF_4_) (99 %), potassium hexacyanoferrate (III) (K_3_[Fe(CN)_6_]) (99 %), pyocyanin (PYO, ≥ 98 %), 2,6-dichloro-1,4-benzoquinone (DCBQ, 98 %) and 3-(3,4-dichlorophenyl)-1,1-dimethylurea (DCMU, ≥98 %) were purchased from Sigma-Aldrich. 3, 4-Ethylenedioxythiophene (EDOT) (> 98 %) was purchased from TCI. Hydrogen peroxide (30 %) (H_2_O_2_) and ammonium hydroxide (25 %) (NH_4_OH) were purchased from Analer Normapur and J.T. Baker, respectively. n-Butanol (AR grade) and pyridine (AR grade) were obtained from Lab-scan analytical sciences. Acetonitrile (MeCN) (HPLC-isocratic grade) was obtained from Hipersolv Chromanorm. All chemicals were commercially available and used without further purification and all solutions were prepared with deionized Milli-*Q* water. TBABF_4_ was dried in vacuum oven at 75 °C for 2 h before use. MeCN was stored over molecular sieves (4 Å, Sigma-Aldrich) for more than 24 h prior to use.

#### 2.1.2. Cyanobacterial cell culture

We used the wild-type cyanobacterium *Synechocystis* sp. PCC 6803 (hereafter *Synechocystis*). Cultures were grown in 30 ml batches of BG11 medium (pH 8.2, 10 mM TES-KOH) [65] at 30°C under continuous white light of 50 µmol_photons_ m^−2^ sec^−1^ under atmospheric CO_2_ with shaking 120 rpm. The components of the BG11 medium were: 17.6 mM NaNO_3_, 0.3 mM MgSO_4_, 245 µM CaCl_2_, 188.7 µM Na_2_CO_3_, 172.2 µM K_2_HPO_4_, 46.3 µM boric acid, 31 µM citric acid, 12.3 µM ferric ammonium citrate, 9.1 µM Mn Cl_2_, 2.7 µM EDTA-Na_2_, 1.9 µM Na_2_MoO_4_, 771.9 nM ZnSO_4_, 316.4 nM CuSO_4_, 171.2 nM Co(NO_3_)_2_.[65] At the logarithmic growth phase, cells were harvested and resuspended in fresh BG11 (pH 8.2) at OD_750_ of 0.1 for sub-culture. Cultures of early stationary phase cells at OD_750_ of ca 1 were concentrated by centrifugation at 6000 *g* for 20 min, the supernatant removed, and the pellet resuspended in fresh BG11 medium pH 8.2 to a chlorophyll *a* (Chl) concentration of 150 nmol_Chl_ ml^−1^ to be used for experiments. Culture chlorophyll concentration was calculated from absorbances at 680 and 750 nm: (A_680_-A_750_)×10.814.[66] All measurements were obtained using a UV-1800 spectrophotometer (Shimadzu).

#### 2.1.3. Electrode fabrication

AP-VPP PEDOT electrodes were fabricated as described previously and illustrated in Figure 1B.[21] In brief, substrates such as glass and PET films were cleaned by ultrasonication in acetone, water, and ethanol. Prior to oxidant solution spin coating, substrates were oxygen plasma treated. The oxidant solution of 0.236 M FeTOS containing 0.141 M pyridine was prepared in n-butanol and spin-coated at 2400 rpm for 80 s. Solvent traces in the spin-coated oxidant thin film were removed by heating at 90 °C for 90 s. The oxidant-coated substrate was immediately transferred to the AP-VPP cell containing EDOT monomer at 75°C. After the polymerization for 4 min, the oxidant-coated substrate comprising PEDOT film was annealed at 90°C for 90 s and thoroughly dip-rinsed in ethanol, followed by acetonitrile. The washed PEDOT film was dried under a dry nitrogen gas stream and stored in a desiccator before use/application. The procedure was repeated from the oxidant spin coating step on PEDOT films to produce multilayered PEDOT films.

### 2.2. Characterization

#### 2.2.1. Sheet resistance and conductivity

Jandel RM3000+ test unit equipped with a Jandel multi-height 4 points cylindrical probe head with load 60+, a tip radius of 100 μm and spacing of 1 mm was used to determine the sheet resistance (ρ_sheet_) of the films. The ρ_sheet_ values were recorded as the average of three measurements.

#### 2.2.2. Electrochemistry and *in situ* spectroelectrochemistry

Cyclic voltammetry experiments were conducted using Metrohm Autolab PGSTAT 101 potentiostat or IviumStat potentiostat/galvanostat instrument (Ivium Technologies) in a conventional three-electrode configuration using a one-compartment Teflon cell. AP-VPP synthesized one layer (1L) and two layer (2L) PEDOT films were used as working electrodes, along with an Ag/AgCl wire as a reference electrode, and a coiled platinum wire as a counter electrode. The reference electrode was calibrated for every electrochemical measurement using the K_3_[Fe(CN)_6_] redox couple (E_1/2_ (Fe/Fe^+^) = 0.248 V). All potentials are given versus this reference electrode. The area of the working electrode was 0.95 cm^2^. All the cyclic voltammograms (CVs) were recorded in 0.01, 0.05 and 0.1 M KCl or NaNO_3_ or in BG11 using scan rates of 20, 50, 100, and 150 mV/s as specified in the figure captions in a potential range of −0.25 to 0.75 V. The electrolyte solution was purged with dry nitrogen gas for 15 min prior to measurement. The charge (Q) was calculated by integration of the cyclic voltammogram in the potential range −0.25 V to 0.75 V.

For the *in situ* UV–Vis spectroscopy measurements, the AP-VPP PEDOT film was used as working electrode in a 1 cm path length quartz cuvette. An Ag/AgCl wire and platinum wire were used as reference and counter electrodes, respectively, and 0.1 M TBABF_4_-MeCN was used as electrolyte solution. The UV-Vis spectra were recorded between 300 and 1100 nm using an Agilent Technology Cary 60 UV-Vis spectrometer. The spectra were recorded in the potential range from –0.9 V to 0.8 V using 100 mV steps.

#### 2.2.3. Scanning electrochemical microscopy

Cyanobacterial cells were loaded onto PEDOT electrodes following the bio-photoelectrochemistry experiment protocol (section 2.2.5.). After gently removing the electrolyte from the photoelectrochemical cell, the loaded electrode was taken out and air-dried for 24 hours, then subjected to further drying in a plasma cleaner under vacuum. The electrode was then stored in a desiccator until imaging (**Supplementary Figure 4A**). The loaded electrodes were mounted on aluminium stubs with copper tape (**Supplementary Figure 4B**) and coated with 10 nm platinum using a Quorum Q150V ES+ sputter coater (**Supplementary Figure 4C**). The sample was imaged using a Apreo Scanning Electron Microscope (Thermo Fischer) with a 2 kV beam acceleration.

#### 2.2.5. Biocompatibility assessment through photosynthesis and cell growth

Biocompatibility was tested by measuring the photosynthetic activity and cell growth when exposed to PEDOT electrodes or glass (only substrate as the negative control) by the methods previously reported.[67] In brief, 3 ml of a cell suspension (OD_750_ = 1.0) was drop-cast onto the substrates placed in a Petri dish. The cells were incubated for 24 h under culture conditions after which the photosynthetic activity and growth of the cells was measured. The photosynthetic yield of photosystem II (Y(II)), was determined by measuring room temperature Chl fluorescence of the cells on the substrates after 10 minutes dark adaptation using a pulse-amplitude modulation fluorimeter (DUAL-PAM-100, Walz, Germany). The cells were illuminated with a red light-saturating pulse (6000 µmol photons m^−2^ s^−1^, 500 ms) to measure maximum fluorescence. The cell growth was measured on solid medium (i.e. a spot test). Cells were harvested from the substrate and resuspended in fresh BG11 (pH 8.2) at OD_750_ = 0.5. Aliquots (10 μl) of three serial dilutions (1×, 10^−3^ and 10^−6^) were spotted onto BG11 agar (pH 8.2) and incubated under culture conditions (section 2.1.2.) for 7 days. Photographs were taken of the plates and growth assessed. The biocompatibility experiments were performed in triplicate.

#### 2.2.5. Bio-photoelectrochemistry

The photoelectrochemical measurements were performed in a three-electrode electrochemical cell (**Supplementary Figure 1**). The AP-VPP-fabricated PEDOT electrodes or ITO-coated glass wafers (Delta Technologies) were placed on the floor of the single chamber cell with a coiled platinum counter electrode and an Ag/AgCl wire reference electrode inserted in the chamber from above. All photoelectrochemistry measurements were recorded under atmospheric conditions at approximately 22 ± 2°C.

150 μl of the 150 nmol_Chl_ ml^−1^ cell concentrate was drop cast onto the working electrode inside the electrochemical cell. 1.35 ml of BG11 (pH 8.2) electrolyte was gently added on top. All experiments were performed in this total 1.5 ml BG11 (pH 8.2) electrolyte solution. The opening of the electrochemical cell was covered with parafilm to reduce evaporation of the electrolyte. The loaded electrodes were left in the dark for 16 h at room temperature before analysis.

Chronoamperometry experiments without a mediator were performed at the applied potential stated in the figure caption (Fig. 5-8). Chronoamperometry experiments with exogenous electron mediators 500 µM pyocyanin (PYO), 1 mM 2,6-dichloro-1,4-benzoquinone (DCBQ) and 1 mM potassium ferricyanide (K_3_Fe(CN)_6_) were performed at applied potentials of 0.1, 0.3 and 0.5 V vs Ag/AgCl, respectively.

During chronoamperometry, the cells were illuminated from below through the electrode with a ModuLight programmable light source (the add-on module operated in combination with the IviumStat, Ivium Technologies) using a wavelength and light intensity as specified in the figure captions (Fig. 5-8, Supplementary Table 2). Photocurrent profiles were measured under 2 min/2 min light/dark cycles; the sixth photocurrent profile was taken for analysis for no exogenous electron mediator, and the first photocurrent profile was taken for analysis for exogenous electron mediator. For longevity experiments, 1 min/59 min light/dark cycles were used.

The photocurrent outputs were calculated as the difference in steady-state current output in the last 5 seconds of the light and dark periods. The calculated photocurrent outputs were normalized to the nominal exposure area of the electrode (0.95 cm^2^) that was covered with cells and exposed to electrolyte and light to obtain photocurrent densities. For exogenous electron mediator experiments, the photocurrent enhancement with exogenous mediator added was calculated relative to the photocurrent without an exogenous mediator added. For longevity experiments, the photocurrent at each hour was calculated as a percentage relative to the initial photocurrent.

#### 2.2.6. Electrode bending

The initial sheet resistance of AP-VPP PEDOT and ITO films on polyethylene terephthalate (PET) substrates were determined by the mean value of at least three different measurements with the same conditions by using a Jandel four-point probe system. After that the substrates were bent using a lab-made bending system with a curvature diameter of 15 mm. At each number of bending cycles (50, 150, 350, 600 and 1200) the movement was paused to measure the sheet resistance values. The values were measured from the square-shaped samples from four corners, relatively far from the strained region. Also the influence of curvature direction was checked. PET films with a thickness of 125 μm were used as flexible substrates. PET and PET-ITO sheets were purchased from GoodFellow and Sigma-Aldrich, respectively.

## 3. Results and discussion

### 3.1. Electrochemical and microscopic characterisation of cyanobacterial cell-AP-VPP fabricated PEDOT electrode

Single layer PEDOT films were fabricated onto flat, non-conductive cleaned glass substrates using the AP-VPP technique that was previously optimized.[21] While the electrochemical performance of AP-VPP fabricated PEDOT films had been previously analyzed using cyclic voltammetry in organic electrolyte (0.1 M tetrabutylammonium tetrafluoroborate in acetonitrile),[21] in this study, their electrochemical performance was further analysed in aqueous electrolytes. In a three electrode photoelectrochemical cell (**Supplementary Figure 1**), we explored KCl, NaNO_3_ and BG11 electrolytes. The latter serves as the medium for the cultivation of the freshwater cyanobacterium *Synechocystis*.[65]

The AP-VPP-fabricated PEDOT electrodes demonstrated electrochemical activity in the aqueous electrolytes, including BG11, as indicated by the CVs (**Figure 2A**), making them suitable for interfacing cyanobacterial cells and further exploration of aqueous electrolytes. To explore the role of diffusion-controlled processes in these aqueous electrolytes, cyclic voltammograms were conducted at different scan rates (**Supplementary Figure 2A-G**). Charge densities were calculated from these CVs and subjected to linear regression against the scan rate (**Figure 2B**). The solid lines in the same graph are the linear fitting of the data. The trends in the slopes of these lines for each electrolyte are shown in **Supplementary Figure 3**. A negative linear correlation between charge density and scan rate indicates the prevalence of diffusion-controlled processes.

**Figure 2.**
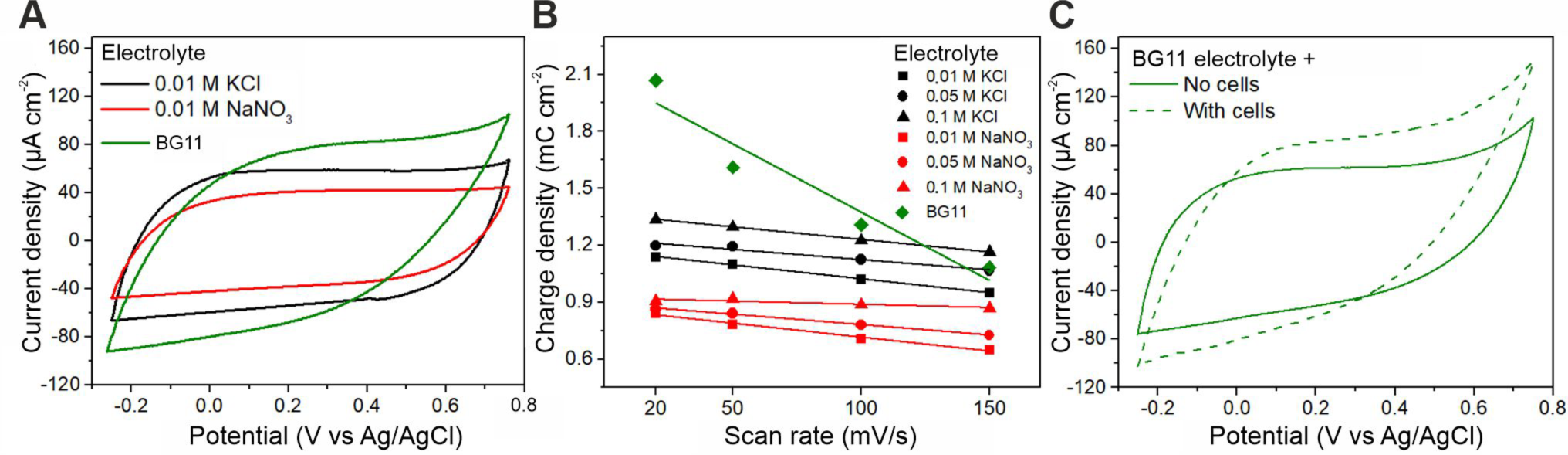
Electrochemical performance of AP-VPP-fabricated PEDOT electrodes. **A)** Cyclic voltammograms in 0.01 M KCl (black line), 0.01 M NaNO3 (red line), and BG11 (green line) electrolytes. **B)** Comparison of charge densities (calculated from the CVs in Supplementary Figure 2 A-G when using 0.01, 0.05, and 0.1 M concentrations (different symbols) of KCl (black) and NaNO3 (red), and BG11 (green diamond) electrolytes. Solid lines are linear fits, the gradients of these lines are given in Supplementary Figure 3. **C)** Cyclic voltammograms in BG11 (pH 8.2) electrolyte without (green solid line) and with *Synechocystis* cyanobacterial cells loaded (green dotted line). Photoelectrochemical parameters: single layer AP-VPP-fabricated PEDOT electrodes, 20 nmolChl cell loading in panel C with cells, cyclic voltammetry at scan rate 100 mV s^-1^ in panels A and C, darkness. One film of each shown.

The negative linear correlation between charge density and scan rate remained relatively constant for KCl electrolytes at different concentrations (**Figure 2B, Supplementary Figure 3**), showing no effect of KCl concentration change on diffusion-controlled doping-dedoping processes. In contrast, the negative linear correlation between charge density and scan rate diminished with increasing NaNO_3_ concentration (**Figure 2B, Supplementary Figure 3**), indicating the effect of increased ionic conductivity. At 25 °C, the difference in the diffusion coefficient of Cl^-^ (2.03 × 10^-9^ m^2^s^-1^) and NO_3_^-^ (1.90 × 10^-9^ m^2^s^-1^), and K^+^ (1.96 × 10^-9^ m^2^s^-1^) and Na^+^ (1.33 × 10^-9^ m^2^s^-1^) can explain the enhanced effect of ionic conductivity of relatively slow diffusing NaNO_3_.[68] Furthermore, at concentrations lower than 0.1 M, the electrical conductivity of the KCl aqueous solution is 52-48 % higher than that of NaNO_3,_ and indeed increased charge densities were observed in KCl electrolytes compared to NaNO_3_ electrolytes (**Figure 2B**). These results demonstrate the importance of ion composition in the electrolyte for high performance of AP-VPP-PEDOT electrodes.

In BG11 medium, 0.0176 M NaNO_3_ is the substantial contributor to ionic strength as the other ionic components of BG11 medium are in micromolar amounts.[65] The negative linear correlation between charge density and scan rate is 5 to 7-fold higher for BG11 compared to NaNO_3_ electrolytes (**Supplementary Figure 3**). This is evidence of a matrix effect with BG11 (i.e. resistance due to bulky, slow diffusing ions/molecules/compounds). Similarly, the deteriorating shape of capacitive current in the CV measured in BG11 electrolyte (**Figure 2A**) is due to this matrix resistance to diffusing anions. The further enhanced matrix resistance can be observed in the CVs of AP-VPP fabricated PEDOT electrodes loaded with *Synechocystis* cyanobacterial cells in BG11 electrolyte (**Figure 2C**). The high charge density in BG11 electrolyte at a low scan rate (20 mV/s) (**Figure 2B**, **Supplementary Figure 2G**) signifies that the maximum doping in the PEDOT electrode occurred in the BG11 matrix compared to NaNO_3_ and KCl electrolyte solutions.

A future approach to boosting the outputs from BPVs using PEDOT electrodes could be to use non-model photosynthetic microorganisms that are tolerant of high levels of salt and different ions, such as estuarine environment strains like *Synechococcus* sp. PCC 11901.[69,70] These strains would enable use of electrolytes with high ionic concentrations and ion compositions suitable for high PEDOT performance, without compromising the photosynthetic and therefore exoelectrogenic performance of the biocatalysts. Although further characterisation would be needed, as it is known that CPs have strain-specific interactions.[56] *Synechocystis*, BG11 was used as the electrolyte solution in subsequent photoelectrochemical experiments.

To examine the arrangement of cells on the electrode that may account for the differences in electrochemical performance (**Figure 2C**), cell-loaded AP-VPP-fabricated PEDOT electrodes (**Figure 3A**) were removed carefully from the photoelectrochemical cell, gently washed with BG11 to remove loosely adherent cells, and visualized by scanning electron microscopy (SEM). The cyanobacterial cells were observed to pack tightly together and interface with the AP-VPP-fabricated PEDOT electrodes in a uniform monolayer (**Figure 3B**), which extended uniformly to the edge of the O-ring in the photoelectrochemical cell (**Figure 3C**). Nevertheless, the micrometer-sized spaces and cavities visible in the SEM images (**Figure 3B,C)** must significantly facilitate the transport of ions to/from the electrode surface as the capacitive currents in the CVs with/without cell loading did not change considerably (**Figure 2**), except for deterioration caused by the cell-matrix effect.

**Figure 3.**
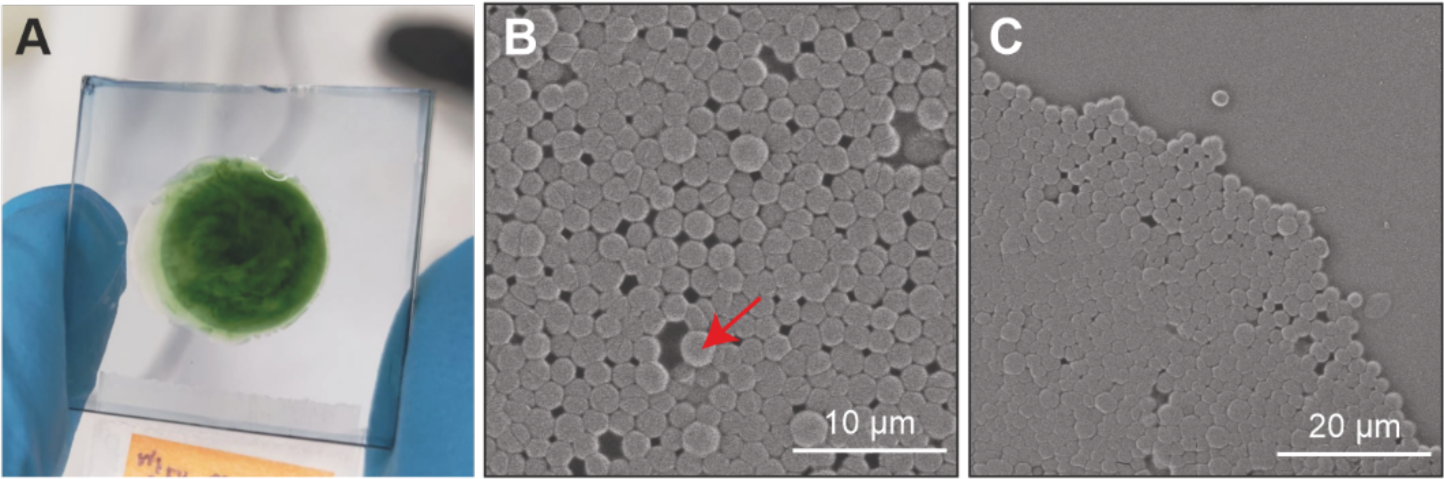
Visualising cyanobacterial cells interfacing with AP-VPP-fabricated PEDOT electrodes. **A)** Photo of a cell-loaded AP-VPP-fabricated PEDOT electrode after photoelectrochemistry experiments. Scanning electron microscope images of **B)** the middle of the cell-loaded electrode and **C)** the edge where the O-ring was. Red arrow – cell displaced from the monolayer.

From the SEM images one can observe that some cells were displaced from the monolayer (**Figure 3B**, red arrow), which could be attributed to the sample preparation process for microscopy. The displaced cells sometimes revealed cracks in the electrode below (**Supplementary Figure 5A**). Additionally, extracellular matrix of exopolysaccharides and some type IV pili connecting the cells on the surface of the PEDOT were also observed (**Supplementary Figure 5B,C**). Lastly, the occurrence of cell division within the biofilm formed on the electrode was observed (**Supplementary Figure 5D**).

### 3.2. AP-VPP-PEDOT electrodes are biocompatible with photosynthesis and growth

The biocompatibility of the AP-VPP-fabricated PEDOT electrodes was tested by measuring the photosynthetic activity and cell growth of *Synechocystis* cyanobacterial cells after 24 h exposure to the electrodes. As a control, the cells were also exposed to a cleaned glass wafer, which served as the substrate for the fabrication of the AP-VPP PEDOT electrodes.

To assess the photosynthetic activity of the cells interacting with the materials, the change in the photosynthetic yield of photosystem II in a dark-adapted state, Y(II), was measured. This is a common indicator of photosynthetic activity and is sensitive to environmental stress.[71] The *Synechocystis* cells on the PEDOT electrodes exhibited typical Y(II) values (0.39 ± 0.003), similar to cells on glass only (0.38 ± 0.026, P = 0.6704, n = 3) (**Figure 4A**). This indicates that the AP-VPP-fabricated PEDOT electrodes maintain the photosynthetic activity of the *Synechocystis* cyanobacterial cells. Consequently, they are suitable as biocompatible anodes for BPV devices, where current is harvested from photosynthesis and an intimate cell contact and immobilization is needed.

**Figure 4.**
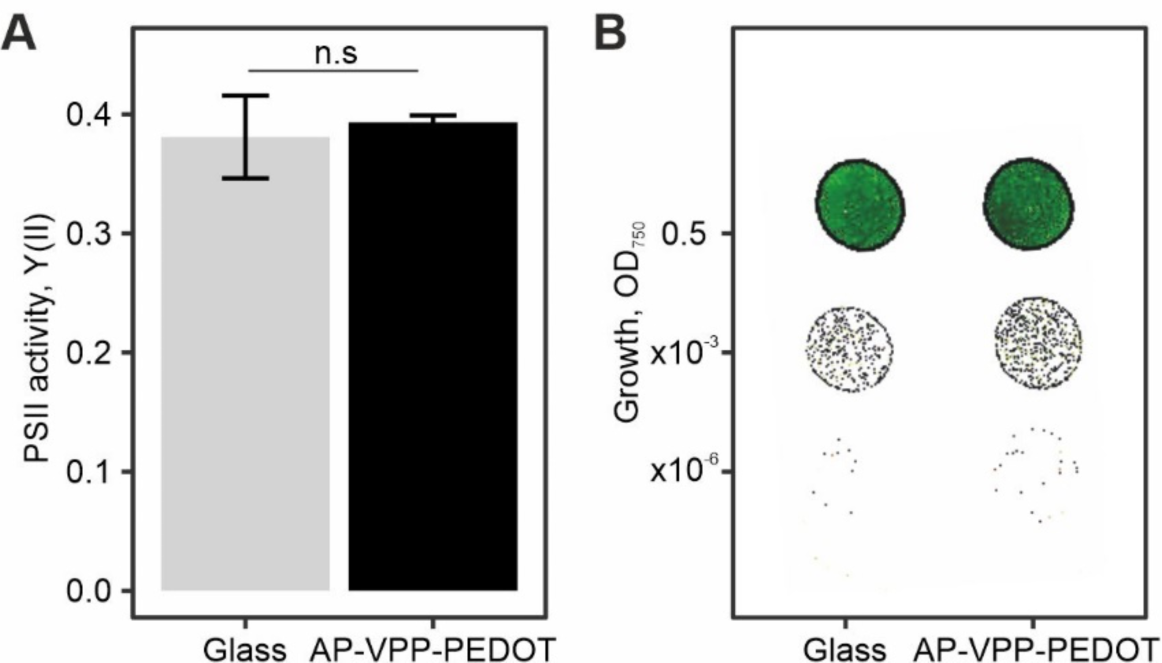
Characterisation of VPP-PEDOT as biocompatible material for interfacing with cyanobacterial cells as a bio-anode in biophotovoltaic devices. **A)** Photosynthetic yield, Y(II), of cells 24 h after loading on electrodes. Data presented as the mean ± SEM of 3 biological replicates, statistical non-significance by t-test P > 0.05 denoted by n.s. **B)** Cell growth on BG11 agar for 7 days following 24 h exposure on electrodes. Glass with no PEDOT coating was used as the negative control. Raw photos of all replicates in (**Supplementary Figure 6**).

The evaluation of cell growth after exposure to the materials on solid BG11 medium with agar (pH 8.2) demonstrated that cells exposed to both the PEDOT electrodes and the glass substrates exhibited similar growth (**Figure 4B**). This demonstrates that the AP-VPP-fabricated PEDOT electrodes exhibit good biocompatibility and do not induce cytotoxicity in *Synechocystis* cyanobacterial cells during prolonged exposure (24 h). As a result, they are suitable for long-term bioelectrochemical applications that use photosynthetic microorganisms as living biocatalysts.

### 3.3. Photoresponse of *Synechocystis* on AP-VPP PEDOT electrodes

To assess the photocurrent output of *Synechocystis* cyanobacterial cells on AP-VPP-fabricated PEDOT electrodes in BG11 electrolyte, chronoamperometry experiments under light/dark cycles were conducted. Chronoamperometry was performed at an applied potential of +0.1 V vs Ag/AgCl wire pseudo reference electrode. The Ag/AgCl reference electrode was calibrated using K_3_[Fe(CN)_6_] redox couple (E_1/2_ (Fe/Fe^+^) = 0.248 V). The choice of applied potential was based on a previous study where +0.1 V vs Ag/AgCl (saturated KCl) was found to be the applied potential at which the maximum photocurrent is reached for *Synechocystis* cells (although this study used inverse-opal structured electrodes made of indium tin oxide (IO-ITO)).[72] This is also an applied potential at which the PEDOT is electrochemically active and shows good conductivity (**Figure 2**) and is near the open circuit potential of the cell-free electrodes (150-200 mV vs Ag/AgCl). During chronoamperometry the cell-loaded electrodes were exposed to darkness followed by 460 nm light at 200 µmol photons m^−2^ s^−1^ (approximately 4 mW cm^−2^ equivalent) in 2 min/2 min cycles (during which periods the steady-state currents in the light and dark had stabilized). The currents were normalized to the surface area of the cell-loaded electrode exposed to illumination, yielding current densities.

*Synechocystis* cyanobacterial cells on AP-VPP-fabricated PEDOT electrodes under light/dark cycles yielded a photocurrent profile (changing ‘shape’ of the current as it evolves from the cells upon dark-light-dark transitions) (**Figure 5A**). The photocurrent profiles were complex (i.e. did not simply reach a steady-state directly) and showed similar features as previously reported for *Synechocystis* cyanobacterial cells on IO-ITO electrodes[72,73] and carbon-based electrodes,[63,74–76] although with different kinetics. The photocurrent profiles of *Synechocystis* cyanobacterial cells on AP-VPP-fabricated PEDOT electrodes showed a peak at 4 s after illumination (5 s for IO-ITO)[73] and a trough that reached its minimum at 28 s after illumination (12 s for IO-ITO)[73] before climbing to a steady-state current output under illumination that would reach its maximum 110 s after illumination began (second peak at 25 s and steady-state 60 s after illumination for IO-ITO[73]). Cyanobacteria on PEDOT-DS-coated graphite were studied under 30 min/30 min light/dark cycles and steady state was reached ca 15 min after illumination[55]; cyanobacteria on PEDOT-PSS-painted carbon cloth were studied under 12 h/12 h light/dark cycles[54]. The fast response (seconds timescale) that were obtained in this study with AP-VPP PEDOT electrodes is indicative of thin active material on top of inert glass substrate achieved by optimised layer-by-layer approach.

The photocurrent output was calculated by taking the difference between the steady-state currents in the light and dark periods (**Figure 5A**, green arrow). While there was some light-response from the AP-VPP-fabricated PEDOT electrodes in the absence of cells (1.9 ± 3.7 nA cm^-2^, n = 3), the primary source of photocurrent was attributed to cyanobacterial cells (34.3 ± 14.6 nA cm^-2^, n = 3, P = 0.0329) (**Figure 5B**). The photocurrent predominantly originated from water splitting in photosystem II (PSII), as evidenced by the cessation of photocurrent (0.0 ± 4.6 nA cm^-2^, n = 3, P = 0.0178) upon the addition of the specific inhibitor, (3-(3,4-dichlorophenyl)-1,1-dimethylurea) (DCMU), which blocks electron transfer from PSII. It has been previously shown that photosynthetic electrons, originating from water-oxidation, are eventually exported from the cells through exoelectrogenesis.[77]

The effect of cell loading was also investigated, and it was observed that the photocurrent output had reached a maximum by 20 nmol chlorophyll loading as higher cell loadings did not result in enhanced photocurrent output (**Supplementary Figure 7**). After the photoelectrochemistry experiments, the cells that remained adherent on the electrode were harvested and measured the chlorophyll content. Approximately 10% of initially loaded cells remained on the electrode surface at 20 nmol_Chl_ loading (1.86 ± 0.39 nmol_Chl_) (**Supplementary Figure 8**). These findings suggest that the biocatalyst loading is not the limiting factor influencing the photocurrent output under these conditions. Moreover, SEM analysis revealed that the cells were densely packed on the flat electrode interface (**Figure *3*B**), supporting the notion that the maximum cell coverage was achieved when using 20 nmol_Chl_ loading.

To determine the long-term stability of the AP-VPP PEDOT electrodes, which is an important consideration in practical applications as well as for analytical studies, the photocurrent outputs of the *Synechocystis* cyanobacterial cells were measured over a 10 h period. The photocurrent output each hour was calculated relative to the initial photocurrent output. The photocurrent output of the *Synechocystis* cyanobacterial cells did not significantly decrease across the 10 h period (0 h: 84.5 ± 65 nA cm^-2^, 11 h: 47.0 ± 29.7 nA cm^-2^, n = 3, P = 0.6293) (**Figure 5C**). Therefore, the cells were stable on PEDOT over 10 h which matches the longevity of cells on IO-ITO electrodes reported previously.[72,73]

### 3.4. Effect of applied potential and exogenous electron mediators

As a CP, PEDOT can be electrochemically oxidized (p-doped) in a reversible way in the presence of electrolytes. During the electrochemical doping reaction CPs are involved in two kinds of kinetic processes: the charge carrier transport along and between the polymer chains, and the movement of counter ions in the film. The broad shape of the CV (**Figure 2C**) can be explained as the consecutive oxidation of polymer segments with different conjugation lengths. At -0.2 V the film is in its neutral to partially oxidized form and starts to be more conducting (formation of charge carriers) when going towards potentials 0 V and 0.6 V (**Figure 2**). Therefore, the conductivity of the PEDOT electrode changes at different applied potentials, and how this effects its ability to harvest current from cyanobacteria was subsequently investigated. Chronoamperometry experiments were conducted at different potentials (0.1, 0.3 and 0.5 V vs Ag/AgCl) under 2 min/2 min 460 nm light at 200 µmol photons m^−2^ s^−1^ (approximately 4 mW cm^−2^ equivalent)/dark cycles.

Higher applied potentials yielded 13-fold greater photocurrent output (15.8 ± 3.3 nA cm^-2^, n = 4 at 0.1 V; 208.2 ± 16.1 nA cm^-2^, n = 5 at 0.5 V applied potential, P = 0.0001) (**Figure 6A**), which is consistent with the p-doping of the PEDOT. This is different to the performance of IO-ITO, where the maximum photocurrent was achieved at 0.1 V vs Ag/AgCl.[72] The physiological effects of subjecting photosynthetic microorganisms to higher applied potentials needs to be explored. It is important to note that the longevity in this study was performed at 0.1 V vs Ag/AgCl (**Figure 5C**).

**Figure 5.**
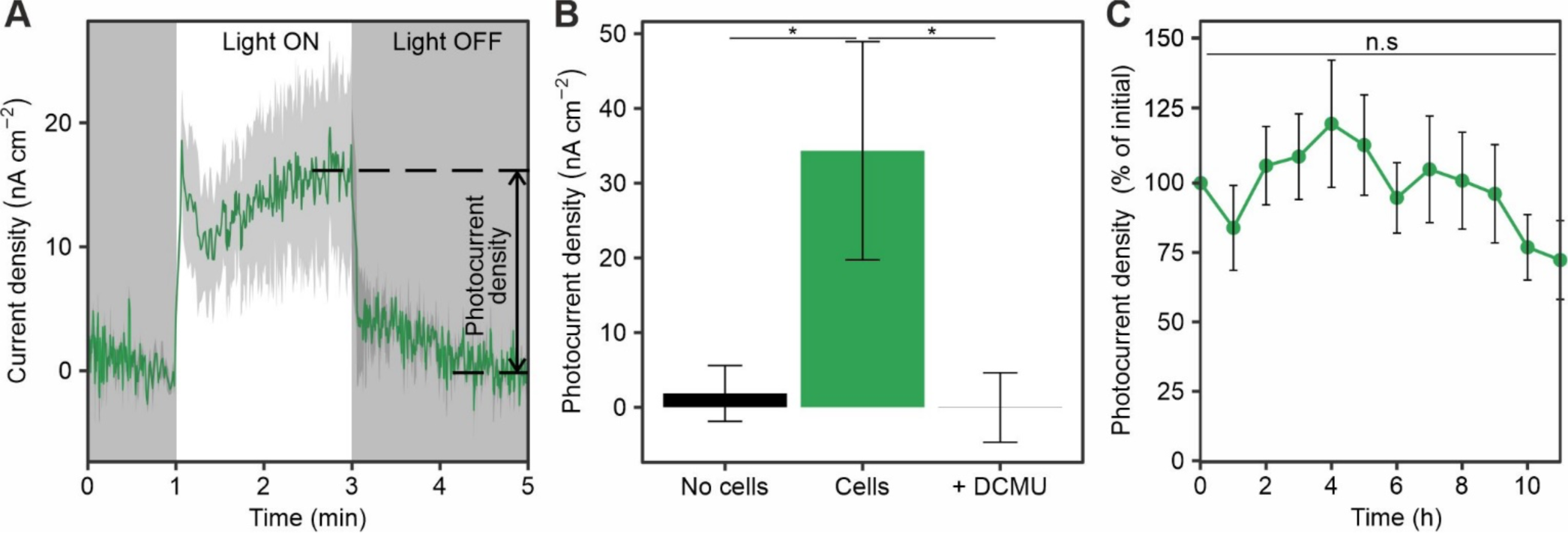
Photoelectrochemical performance of *Synechocystis* cells on AP-VPP-fabricated PEDOT electrodes. **A)** Photocurrent profile, and **B)** Photocurrent outputs, which were calculated as the difference between the steady-state currents in the light and dark periods of the photocurrent profile (arrow in panel A). **C)** Longevity of *Synechocystis*:AP-VPP-PEDOT bioanodes. Photocurrent outputs from *Synechocystis* cyanobacterial cells over 10 h as a percentage of the initial photocurrent output. Photoelectrochemical parameters: single layer AP-VPP-fabricated PEDOT electrodes, 20 nmolChl cell loading, chronoamperometry at 0.1 V vs Ag/AgCl applied potential, 4 mW cm^−2^ 460 nm light, 2 min/2 min light/dark cycles. Data presented as the mean (solid line in photocurrent profile) ± standard error of the mean (grey ribbon in photocurrent profile) of n = 3 biological replicates, statistical significance by t-test P < 0.05 denoted by *, P > 0.05 not significant (n.s.).

**Figure 6.**
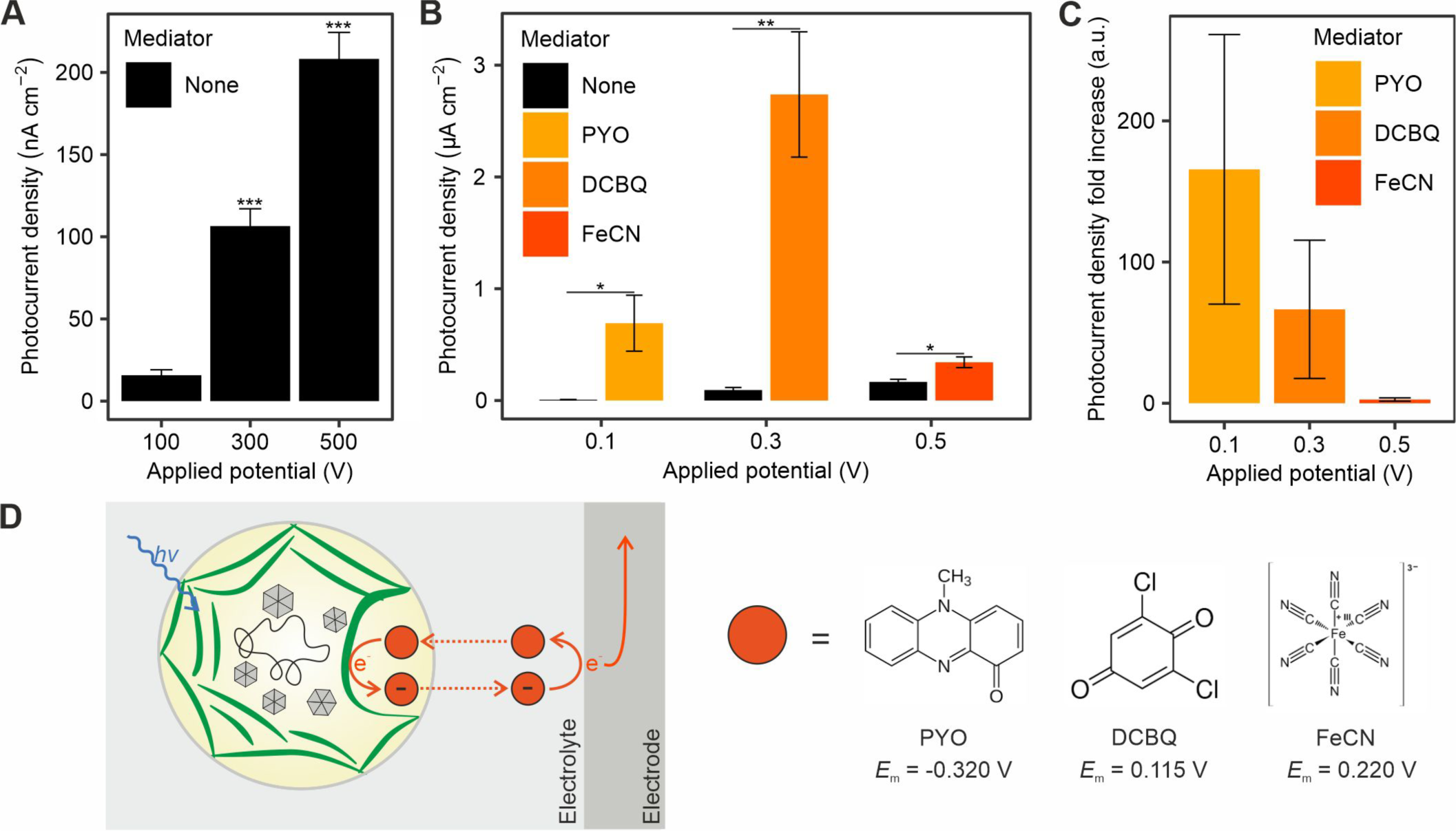
Effect of applied potential and exogenous electron mediators with different mid-point potentials on photoelectrochemical performance of *Synechocystis* cells on AP-VPP-fabricated PEDOT electrodes. **A)** and **B)** Photocurrent outputs of *Synechocystis* cells on AP-VPP-fabricated PEDOT electrodes at different applied potential and exogenous electron mediators added. None – no exogenous electron mediator added, PYO - pyocyanin, DCBQ - 2,6-Dichloro-1,4-benzoquinone, FeCN - potassium ferricyanide. Panels A and B exhibit different scales, nonetheless, no mediator data is identical, facilitating comparison. **C)** Photocurrent outputs from panel B in the presence of different exogenous electron mediators relative to no exogenous electron mediator. No change is 1 arbitrary unit (a.u.). Photoelectrochemical parameters: single layer AP-VPP-fabricated PEDOT electrodes, 20 nmolChl cell loading, chronoamperometry at applied potentials specified vs Ag/AgCl, 4 mW cm^−2^ light at 460 nm (blue), 2 min/2 min light/dark cycles. Data presented as the mean ± standard error of the mean of at least n = 3 biological replicates, statistical significance by t-test P ≤ 0.0001 denoted by ***, P ≤ 0.05 denoted by *. **D)** Schematic of boosting the photocurrent output using exogenous electron mediators. Mid-point potential of employed exogenous electron mediators at pH 7 versus Ag/AgCl.[80,81,83]

Cells naturally have endogenous electron mediator(s) that participate in inefficient indirect extracellular electron transfer,[63,64,72] so boosting how the mediator(s) traverses the cell wall is a point of optimisation.[63,73,78] Artificial or exogenous electron mediators can be used to facilitate extracellular electron transfer, and with different mid-point potentials enabling harvesting of photocurrent at different applied potentials with optimized thermodynamic losses (**Figure 6D**).[79] The photocurrent can be boosted at lower applied potentials using low mid-point potential exogenous electron mediators of the class phenazine, including pyocyanin (PYO), that are naturally produced by a few clades of bacteria, most notably pseudomonads such as *Pseudomonas aeruginosa,* and can also be produced by genetically engineered *Synechocystis*.[80] The photocurrent can be boosted at intermediate applied potentials using a variety of exogenous/artificial quinones,[81] including 2,6-dichloro-1,4-benzoquinone (DCBQ) that harvests electrons from the Q_B_ pocket of PSII as well as from the photoexcited reaction centers of PSII and PSI.[82] At higher potentials, the photocurrent can be boosted using potassium ferricyanide (K_3_[Fe(CN)_6_]), which efficiently harvests electrons from the periplasmic space in the outermost compartment of the cyanobacterial cell and is commonly used in planktonic BPVs.[77] Chronoamperometry experiments were conducted at different potentials under 460 nm light/dark cycles with the addition of different mid-point potential mediators to investigate how photocurrent harvested by the AP-VPP-PEDOT electrodes could be boosted at different applied potentials.

The photocurrent was boosted 166-fold (7.4 ± 0.5 nA cm^-2^ to 0.69 ± 0.25 µA cm^-2^, n = 4, P = 0.0341) by adding 500 µM PYO (maximum solubility in aqueous BG11) at 0.1 V (**Figure 6B,C**). The photocurrent was boosted 67-fold (94 ± 23 nA cm^-2^ to 2.7 ± 0.6 µA cm^-2^, n = 3, P = 0.0092) by adding 1 mM DCBQ at 0.3 V (**Figure 6B,C**). Under these conditions, the highest photocurrent in this study was achieved of 2.7 ± 0.6 µA cm^-2^ (n = 3). (**Figure 6B**). The photocurrent was only doubled (167 ± 23 nA cm^-2^ to 342 ± 48 µA cm^-2^, n = 3, P = 0.0302) at 0.5 V using 1 mM potassium ferricyanide ([Fe(CN)_6_]^3-^, FeCN) (**Figure 6B,C**) despite 90% of cells loaded being not strongly adhered to the electrode (i.e. planktonic) that would benefit from the artificial electron mediator (**Supplementary Figure 8**). We concluded that this was because PEDOT conductivity was already at its maximum at this potential, where the bottleneck could be the electroactive surface area of the flat electrode. Future studies will aim to fabricate AP-VPP-PEDOT electrodes that have similar geometric surface areas but higher electroactive surface areas using 3D architectures.

### 3.5. Harnessing visible light spectrum for cyanobacteria and PEDOT

The effect of light wavelength on the photoelectrochemical performance of *Synechocystis* cyanobacteria cells deposited on AP-VPP-fabricated PEDOT electrode was investigated. While photosynthetic organisms are able to use a wide range of wavelengths of light to perform photosynthesis (photosynthetic active radiation, PAR: 400 - 700 nm), photosynthesis is most efficient within the red range (600–700 nm) and the blue range (425–450 nm).[84] Therefore, blue (440 nm) and deep red (660 nm) lights were tested under light/dark cycles at a physiologically relevant light intensity of 4 mW cm^−2^ in 2 min/2 min cycles. All photoelectrochemistry experiments were performed with the light shining through the PEDOT electrode.

Blue light (150_photons_ m^−2^ sec^−1^, approximately 4 mW cm^−2^ equivalent) yielded higher photocurrent than the deep red light (200 photons m^−2^ sec^−1^, approximately 4 mW cm^−2^ equivalent) (**Figure 7A**). The difference in performance from different lights increased with applied potential (from 0.1 V to 0.3 and 0.5 V), being a three-fold difference and statistically significant at the higher two applied potentials tested (at 0.5 V 440 nm: 208 ± 36 nA, 660 nm: 66.6 ± 25.5 nA, n = 5, P = 0.0124) (**Figure 7A**).

**Figure 7.**
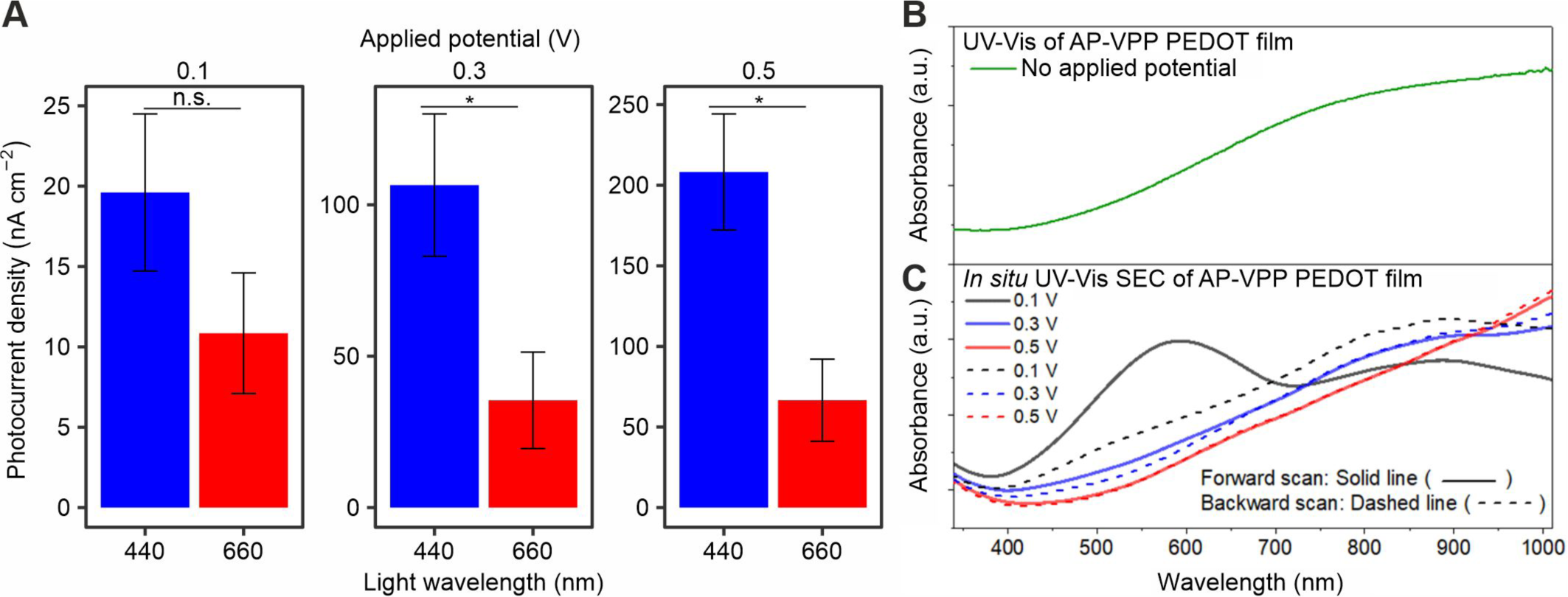
Effect of light wavelength on photoelectrochemical performance of *Synechocystis* cells on AP-VPP-fabricated PEDOT electrodes. **A)** Photocurrent output of *Synechocystis* cells on AP-VPP-fabricated PEDOT electrodes under blue (460 nm) or deep red (660 nm) light. Photoelectrochemical parameters: single layer AP-VPP-fabricated PEDOT electrodes, 20 nmolChl cell loading, chronoamperometry at applied potentials specified vs Ag, 4 mW cm^−2^ light at wavelength as specified, 2 min/2 min light/dark cycles. Data presented as the mean ± standard error of the mean of n = 5 biological replicates, statistical significance by t-test P ≤ 0.05 denoted by *, P > 0.05 not significant (n.s.). **B)** UV-Vis absorbance spectra of AP-VPP-fabricated PEDOT electrode after fabrication not under applied potential, **C)** and under applied potentials during *in situ* cyclic voltammetry. Full *in situ* UV-Vis absorbance spectra coupled with cyclic voltammetry in **Supplementary Figure 9**.

In CPs p-doping.[85] They also cause a reversible change of electrochemical and structural properties such as color, conductivity, porosity, stored charge, or volume. VPP PEDOT is different from other CPs including other members of the polythiophene family as after polymerization (in pristine state) it shows a highly doped/conducting state and a broad absorbance in the Vis range, extending from 430 to 1100 nm (**Figure 7B**). Note that this is the form of AP-VPP PEDOT (with no applied potential) for which the light intensities were calibrated for cell experiments (**Supplementary Table 2, 3**). During light calibration some of the red light shining through the nanometers thick AP-VPP PEDOT film would be absorbed, resulting in a slight underestimation of the red light intensity set.

The change in absorbance of PEDOT at different applied potentials was investigated by conducting *in situ* UV-Vis spectroscopic measurements (SEC). At higher applied potentials, broad-band absorbance between 400 to 650 nm originating from the neutral form of PEDOT shifts to >700 nm as it gets heavily doped, and in conductive form (**Figure 7**, **Supplementary Figure 9**). At 0.1 V applied potential, PEDOT absorbs light mostly in the 500-700 nm range, which includes the red range (**Figure 7**, black line). At this potential the film is in its semi-conducting state and shows two absorption bands centered at 590 and 885 nm. At 0.3 and 0.5 V applied potentials, the absorbance at longer wavelength (above 800 nm) increases at the expense of the absorbance at 590 nm (**Figure 7C**, blue and red lines, respectively) The enhanced absorbance at 590 nm for low oxidation potential is due to the polaronic state formation, while the increase of the absorbance at 885 nm continuing into NIR range for high oxidation potential is due to the bipolaronic state formation.[86,87] Therefore, under the applied positive potential, the steady increase in photocurrent output for both the blue and red light (**Figure 7A**) can be explained by the transformation of PEDOT from a semiconducting to a more conducting form with increased charge carrier formation in the polymer backbone as it oxidizes. Note that for conducting polymers these electro-generated polaronic states are comparatively reversible (**Figure 7C**, dotted lines), although the experiments with cells were done in increasing order of applied potential.

Even though these electro-generated polaronic states have different absorbance spectra, the AP-VPP-fabricated PEDOT electrodes demonstrate unchanged electrochemical activity as measured by cyclic voltammetry under continuous illumination with a white (4100 K) light and six different wavelengths in the visible spectrum at their maximal intensity: blue (460 nm), green (523 nm), amber (590 nm), red (623 nm), deep red (660 nm), far red (740 nm) light (**Supplementary Figure 10**). Furthermore, there was negligible photoresponse from AP-VPP PEDOT at potential of 0.1 V with blue light when no cells were loaded (**Figure 5B**).

The effect of wavelength of light on exoelectrogenesis has been explored in previous studies. The electrogenic (peak power output of a BPV with an ITO-coated PET anode) and photosynthetic activity (oxygen evolution) of green alga *C. vulgaris* were both high under red and blue lights.[88] However, the cyanobacterium *Synechococcus elongatus* exhibited higher electrogenic and photosynthetic activity under red light compared to blue light. Several studies have noted that cyanobacteria with phycobilisome-based light-harvesting antennae use blue light less efficiently for photosynthesis than other photosynthetic organisms with chlorophyll-based light-harvesting antennae, such as green algae. Exposure to blue light in cyanobacteria causes an imbalance between the two photosystems, characterised by a higher association of phycobilisomes with PSII and a lower ratio of PSI to PSII protein complexes.[89] Another study found that the electrogenic activity (change in potential of a BPV with an external resistor with carbon paint anodes coated with the conducting polymer polypyrrole) of sheathed cyanobacteria *Nostoc* and *Lyngby* was similar under red and blue lights.[52]

In this study, blue light yielded higher photocurrent than red light (**Figure 7A**). Therefore, we concluded that the wavelength of light, together with the applied potential, effected PEDOT’s capacity to harvest current from *Synechocystis* cyanobacteria in unexpected ways. This calls for further exploration, including simultaneous studies of electrode properties, photosynthesis and exoelectrogenesis, as well as optimisation in future studies. A wavelength shifting strategy could be employed to optimise light wavelengths for both the PEDOT and the *Synechocystis* cells. For example, an EDOT monomer unit could be synthesized with a fluorene unit that could incorporate nanoparticles into the conducting network capable of absorbing and reemitting some light back that the cyanobacteria could utilize.[90]

### 3.6. Comparative analysis of AP-VPP-PEDOT and other electrodes: performance and potential for industrial application and scalability

The performance of AP-VPP-fabricated PEDOT electrodes was compared with commercially available ITO electrodes for harvesting photocurrent from *Synechocystis* cyanobacteria cells. Both electrode types had a flat architecture. The ITO electrodes had a nominal thickness of 125 nm nanoparticle coating on quartz and a surface resistance of 10-20 Ω/sq. In contrast, our AP-VPP-fabricated PEDOT electrodes featured a thinner single-layer of PEDOT coating on glass of 20-30 nm (5-fold thinner than the ITO electrodes) and 250-450 Ω/sq surface resistance (approximately 25-fold more resistant than the ITO electrodes). The *Synechocystis* cyanobacteria cells yielded a higher photocurrent on AP-VPP-fabricated PEDOT electrodes (166.7 ± 40.2 nA, n = 3) than on ITO electrodes (76.3 ± 32.9 nA, n = 3) (**Figure 8A**). While the difference was not statistically significant (P = 0.1529), one would have expected a much higher photocurrent for ITO compared to VP-APP-PEDOT electrodes due to the difference in thickness and surface resistance.

**Figure 8.**
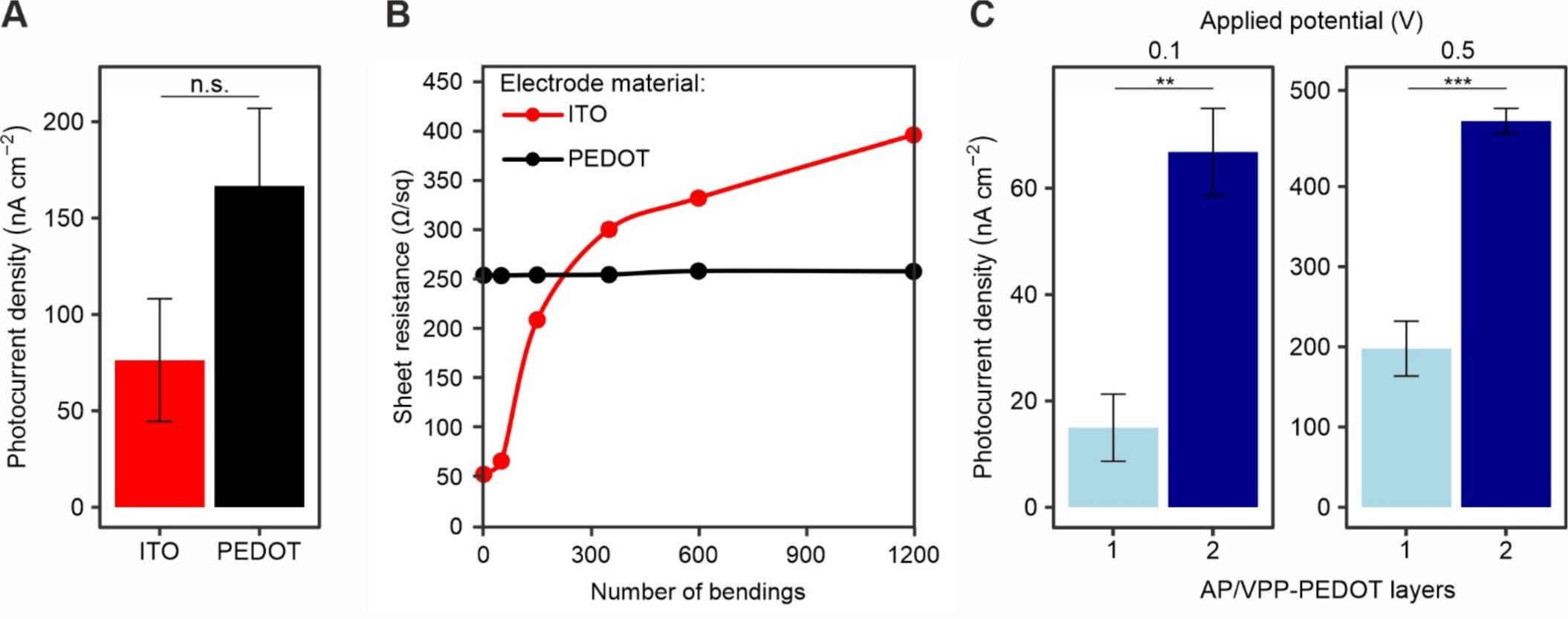
(Photo)electrochemical performance of flat electrodes made of different materials. **A)** *Synechocystis* cells on ITO versus AP-VPP PEDOT on glass. Photoelectrochemical parameters: single layer AP-VPP-fabricated PEDOT or ITO electrodes on flat nonconducting substrate, 20 nmolChl cell loading, chronoamperometry at 0.5 V vs Ag, 4 mW cm^−2^ 460 nm light, 2 min/2 min light/dark cycles. Data presented as the mean ± standard error of the mean of n = 3 biological replicates, P > 0.05 not significant (n.s.). **B)** Sheet resistance measurements of ITO versus AP-VPP PEDOT on PET before and after indicated number of bendings. No *Synechocystis* cells were loaded. Data presented as the mean ± standard error of the mean of forward and reverse measurements, full data in **Error! Reference source not found.**. **C)** *Synechocystis* cells on PEDOT of different layer thicknesses. Photoelectrochemical parameters: 20 nmolChl cell loading, chronoamperometry at applied potentials specified vs Ag, 4 mW cm^−2^ 460 nm light wavelength, 2 min/2 min light/dark cycles. Data presented as the mean ± standard error of the mean of n = 5 biological replicates for one-layer and n = 3 biological replicates for two-layer, statistical significance by t-test P ≤ 0.01 denoted by **, P ≤ 0.001 denoted by ***.

The structural durability of the AP-VPP-fabricated PEDOT electrodes was compared to commercially available patterned ITO electrodes on flexible PET substrate. The electrodes were repeatedly bent with a moderate bending diameter of 15 mm, and their sheet resistance was measured. AP-VPP-PEDOT demonstrated no changes in resistance, whereas ITO exhibited a 7-fold increase in resistance (i.e. decrease in conductivity) after up to 1200 bendings (**Figure 8B**). While an improvement in mechanical flexibility has been reported for ITO nanoarrays or spring-like ITO electrode structures,[91,92] continuous ITO films are fragile and susceptible to crack formation under repeated bending. Previously, continuous ITO-PET substrates failed after a single bending cycle with a bending radius lower than 20 mm.[23] On the other hand, our AP-VPP-fabricated PEDOT films demonstrate flexibility when applied to appropriate substrates like PET or paper. These findings highlight the superior mechanical resilience and conductivity retention of CP PEDOT films compared to ITO films under bending stresses. Additionally, it is noteworthy to emphasise that that the VPP process is conducted under atmospheric pressure (AP), making it an easy and cost-effective way of manufacturing films that can be integrated into roll-to-roll manufacturing processes.

The effect of the thickness of AP-VPP-fabricated PEDOT electrodes on the photoelectrochemical performance of *Synechocystis* cyanobacteria was investigated. The performance of one-layer and two-layer AP-VPP-fabricated PEDOT electrodes was compared using chronoamperometry under light/dark 2 min/2 min cycles of blue (460 nm) light at an intensity of 4 mW cm^−2^ (measured passing through a one-layer PEDOT electrode). Two-layer electrodes yielded approximately 2.5- and 4.5-fold more photocurrent than the one-layer electrodes at 0.1 and 0.5 V applied potentials tested, respectively (at 0.5 V: one-layer 197.8 ± 34.2 nA, n = 5; two-layer 481.6 ± 15.7 nA, n = 3; P = 0.0009) (**Figure 8C**). The photocurrent of two-layer PEDOT at 0.5V and using blue light of 0.46 µA was the highest non-mediated photocurrent achieved in this study. These results demonstrate that PEDOT thickness is an important property for determining photocurrent yield. PEDOT thickness is tunable with the layer-by-layer approach of the AP-VPP fabrication method.

While AP-VPP PEDOT is a promising material, there is a potential for enhancing its single-layer flat architecture into a more advanced hierarchical third-generation electrode.[2] CPs can be printed on existing structures, for example PANI has been inkjet-printed on carbon paper to serve as an anode for non-photosynthetic bacteria in a MFC.[93] In a recent study, 3D-printing techniques have been used to combine PEDOT:PSS and isolated thylakoid membranes.[94] Furthermore, optimising the cell-electrode interface is crucial. This has been achieved with non-photosynthetic organisms by integrating CPs such as PEDOT:PSS and polypyrrole onto the cell walls of *Shewanella oneidensis*, *Aspergillus niger* and *Rhizoctania* sp. in MFCs.[95,96] Recently, polyphenyleneethynylene has been polymerised on the surface of *Chlorella pyrenoidosa* cells.[97] An alternative approach involving a matrix has also been explored; for example *S. oneidensis* was encapsulated in a conductive 3D PEDOT:PSS matrix electropolymerized on a carbon felt substrate, forming a multilayer conductive bacterial-composite film.[98] Future studies employing these strategies with photosynthetic microorganisms in BPVs should carefully consider the effects on light management and photosynthesis.

## 4. Conclusions

In this study, we demonstrated that AP-VPP-fabricated PEDOT can be an environmentally friendly anode material for interfacing with photosynthetic microorganisms in BPVs to generate electricity renewably. We also identified the best parameters to achieve the highest photocurrent outputs in the complicated interplay between the exoelectrogenic photosynthetic cells and light sensitive electropolar electrodes. The highest non-mediated photocurrent achieved was 0.48 µA cm^−2^, which was obtained with a two-layer PEDOT electrode at 0.5 V applied potential using blue light. The highest mediated photocurrent achieved was 2.73 µA cm^−2^, which was obtained with a one-layer PEDOT electrode at 0.3 V applied potential using blue light and the exogenous electron mediator DCBQ. Comparatively, the performances of flat AP-VPP-PEDOT electrodes and flat ITO electrodes were found to be similar, although it should be noted that this was not an ideal comparison due to different film thicknesses and the specific experimental conditions. We achieved a quarter of the present benchmark non-mediated photocurrent outputs of 1.93 µA cm^−2^ (90-fold lower than the benchmark mediated photocurrent output of 245 µA cm^−2^) from electrodes made of ITO and with hierarchical micro-pillar architecture.[7] In future studies we aim to extend the flexible AP-VPP technique to fabricate 3D PEDOT electrodes to achieve higher photocurrent outputs. But also as explored and reported in this study, PEDOT is a dynamic material in different ways, including significantly different photoelectrochemical performance with electrolyte composition, applied potential and exogenous mediators, light wavelength compared to other electrodes. These differences are each unique opportunities for optimisation of the electrode, beyond just optimising architectures in the future to achieve high photocurrent outputs for sustainable green electricity generation.

## CRediT authorship contribution statement

**Laura T. Wey:** Conceptualization, Methodology, Formal analysis, Investigation, Data curation, Writing – Original Draft, Writing – Review & Editing, Visualization, Supervision. **Rahul Yewale:** Methodology, Investigation, Formal analysis, Writing – Original Draft, Writing – Review & Editing, Visualization. **Emilia Hautala:** Investigation, Data curation. **Jenna Hannonen:** Investigation, Data curation. **Kalle Katavisto:** Investigation. **Carita Kvarnström:** Supervision, Resources, Funding acquisition. **Yagut Allahverdiyeva:** Conceptualization, Writing – Review & Editing, Supervision, Resources, Funding acquisition, Project Administration. **Pia Damlin:** Conceptualization, Methodology, Writing – Original Draft, Writing – Review & Editing, Visualization, Supervision, Resources, Project Administration.

## Acknowledgements

We would like to acknowledge Ermei Mäkilä for scanning electron microscopy sample preparation and training, Sergey Kosourov for suggesting the concept of using conducting polymers as an anode and for involvement in preliminary cell studies, Sindhujaa Vajravel for preliminary cell culture and photosynthetic measurements, and Joona Huopalainen for fabrication of AP-VPP PEDOT/PET electrodes. Biophysical experiments were performed within the PHOTOSYN infrastructure at the University of Turku. SmartBio Biocity Turku Research program is acknowledged for stimulating multidisciplinary collaboration.

