## Supplementary information for "Optoelectronic enhancement of photocurrent by cyanobacteria on sustainable AP-VPP-fabricated PEDOT electrodes"

**Supplementary Table 1** Conducting polymers in BPV studies. Carbon nanofiber (CNF), poly(3,4-ethylenedioxythiophene) (PEDOT), polypyrrole (PPy), polyaniline (PANI), polystyrene sulfonate (PSS), dodecyl sulfate (DS), indium tin oxide (ITO).

| **Electrode substrate** | **Conducting polymer** | **Method of polymerisation** | **Photosynthetic microorganism** | **Photocurrent output, comments** | **Reference** |
| --- | --- | --- | --- | --- | --- |
| Carbon cloth, Au layer | PANI | soaking the electrode in the polyaniline solution | *Synechococcus*. sp | Peak current density > 150 µA cm^−2^ with a 100 load, 5.3 µW cm^−2^ of maximum power density | Furukawa et al. 2006[1] |
| Carbon cloth or carbon paint | PANI- or PPy | coated using paint mixed with 10 mg polyaniline emeraldine salt (20 wt.% on carbon black) or 10 mg polypyrrole, un-doped (20 wt.% on carbon black) and air-dry for 10 min, then dried at 70^°^C in a furnace for 5 min | *Synechocystis* sp. PCC 6803 and pond isolate | CP coating gave substantially higher amplitude of the light response compared to bare carbon for pond isolate, not significant improvement for *Synechocystis* | Zou et al. 2009[2] |
| Carbon fibers | PPy | chemical oxidation of pyrrole monomer with ammonium peroxydisulphate | Pond isolate | Loading density of 3 mg/cm^2^, PPy coating resulted in a 450% increase in the power density | Zou et al. 2010[3] |
| Carbon paint | PPy | coated using paint mixed with 10 mg polypyrrole, un-doped (20 wt.% on carbon black) and air-dry for 10 min, then dried at 70^°^C in a furnace for 5 min | Ten genera of cyanobacteria | Electrogenic activity was observed upon illumination | Pisciotta et al. 2010[4] |
| Fluorine doped tin oxide-coated glass | PANI | cyclic voltammetry between 0 and 1.2 V (w.r.t. Ag/AgCl) at a scan rate of 1 mV s^−1^ in a solution of H_2_SO_4_ (1 M) containing 0.1 M aniline | *Pseudanabaena limnetica* | 70.3 ± 0.2 mV in light (55% higher voltage than dark); 54 ± 13 pW (nmol Chl)^−1^ during the light (76 % higher power than dark) | Bombelli et al. 2012[5] |
| Carbon cloth | PEDOT:PSS | brush-painted with a mixture of 1 wt% PEDOT:PSS and 5 wt% DMSO. The DMSO was added to increase the PEDOT : PSS conductivity. All the carbon cloths were exposed to oxygen plasma for 1 min for hydrophilization | Synechocystis sp. PCC 6803 | 438 mW m^−2^ power density output | Liu and Choi 2017[6] |
| Glass-ITO substrate | PEDOT:PSS | PEDOT:PSS-CNF were prepared by mixing PEDOT:PSS/CNF/glycerol/DMSO (final dry weight ratios 16.2/6.9/10.8/77.4). The degassed solution was cast into petri dishes, and dried overnight at 50^°^C | Thylakoid membranes extracted from spinach | 9.0 mC charge on PEDOT:PSS-CNF, compared to 6.6 mC charge on ITO (not significantly different) | Méhes et al 2019[7] |
| Graphite rod | PEDOT-DS | electrosynthesized under 1.3 V potentiostatic chronoamperometric (CA) conditions (100 s) in the presence of a sodium dodecyl sulphate (SDS) as dopant | *Synechocystis* sp. PCC 6803 | CP coating yielded sixfold and twofold increase compared to bare graphite for both mediatorless and ferricyanide-mediated conditions, respectively | Reggente et al. 2020[8] |
| Graphite rod | PPy | polymerization at 1.0 V for 100 s (for 3-mm diameter electrodes) in the presence of a sodium dodecyl sulphate (SDS) as dopant | *Synechocystis sp*. PCC6803 and *Synechococcus elongatus PCC7942* | CP coating yielded sixfold increase in photocurrent for *Synechocystis* under unmediated conditions compared to bare graphite, no significant improvement for *Synechococcus* | Roullier et al., 2023[9] |
| 3D polymeric electrode so no substrate | PEDOT:PSS | PEDOT:PSS polymer matrix dissolved in 15% DMSO solvent to be 6% 3D-printable conductive ink | Thylakoid membranes extracted from spinach | 37.1 μA purely generated from the embedded TMs | Kim et al 2023[10] |

**Supplementary Table 2** Light Intensities used in photoelectrochemistry experiments unless specified otherwise.

| **Electrode material** | **Potential controlling the light (V)** | **Light colour** | **Wavelength (nm)** | **Light intensity (µmol photons m^−2^ s^−1^)** | **Light intensity (mW cm^–2^)** |
| --- | --- | --- | --- | --- | --- |
| Glass | 0.24 | Blue | 460 | 150 |  |
| Glass | 0.39 | Deep Red | 660 | 240 |  |
| Glass | 0.18 | White | NA | 150 |  |
| Single layer PEDOT | 0.24 | Blue | 460 | 150 | 4 |
| Single layer PEDOT | 0.39 | Deep Red | 660 | 200 | 3,75 |
| Single layer PEDOT | 0.18 | White | NA | Not measured |  |

**Supplementary Table 3** Light Intensities used for Supplementary Figure 10.

| **Electrode material** | **Potential controlling the light (V)** | **Light colour** | **Wavelength (nm)** | **Light intensity (µmol photons m^−2^ s^−1^)** | **Power [mW]** |
| --- | --- | --- | --- | --- | --- |
| Single layer PEDOT | 4 | White 4100 K | - | 1800 | 200lm |
| Single layer PEDOT | 4 | Blue | 460 | 1750 | 976 |
| Single layer PEDOT | 4 | Green | 525 | Not measured | 279 |
| Single layer PEDOT | 4 | Amber | 590 | Not measured | 203 |
| Single layer PEDOT | 4 | Red | 623 | Not measured | 461 |
| Single layer PEDOT | 4 | Deep Red | 660 | 2000 | 720 |
| Single layer PEDOT | 4 | Far Red | 740 | Not measured | 405 |

**Supplementary Table 4** Sheet resistance measurements (Ω/sq) of commercial patterned ITO versus AP-VPP PEDOT on PET before (0 bendings) and after (50-1200) number of bendings. No *Synechocystis* cells were loaded. Current feed 1 mA. Data for Figure 8B.

| **Electrode material** | **Number of bendings** | | | | | |
| --- | --- | --- | --- | --- | --- | --- |
|  | **0** | **50** | **150** | **350** | **600** | **1200** |
| **ITO** | 53,9 (FWD)  53,5 (REV) | 67,2  67,2 | 209,8  209,7 | 305,3  297,2 | 336,8  329,5 | 398,4  395,7 |
| **PEDOT** | 254,8 (FWD)  255,2 (REV) | 254,8  254,5 | 254,9  255,4 | 255,3  255,8 | 259,5  258,8 | 259,6  258,1 |

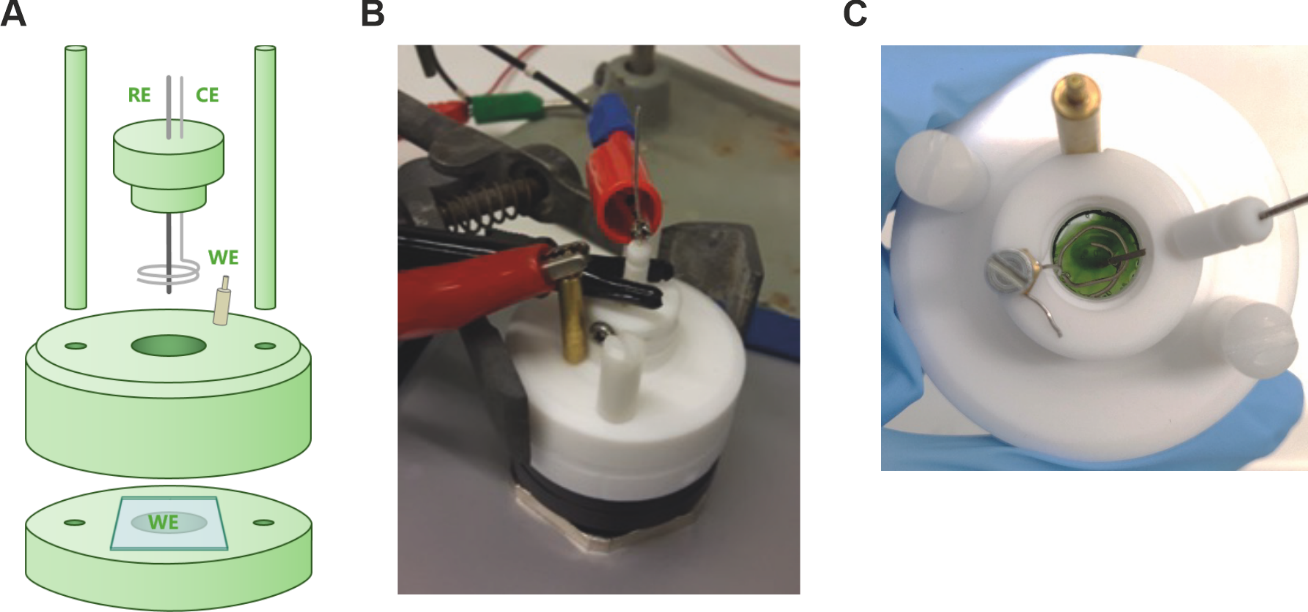

**Supplementary Figure 1** The photoelectrochemical cell used in this study. **A)** Schematic of the photoelectrochemical cell. WE- working electrode, CE - counter electrode, RE - reference electrode. **B)** Photo of the photoelectrochemical cell. **C)** Photo of the cyanobacterial-cell loaded anode in the apparatus viewed from above.

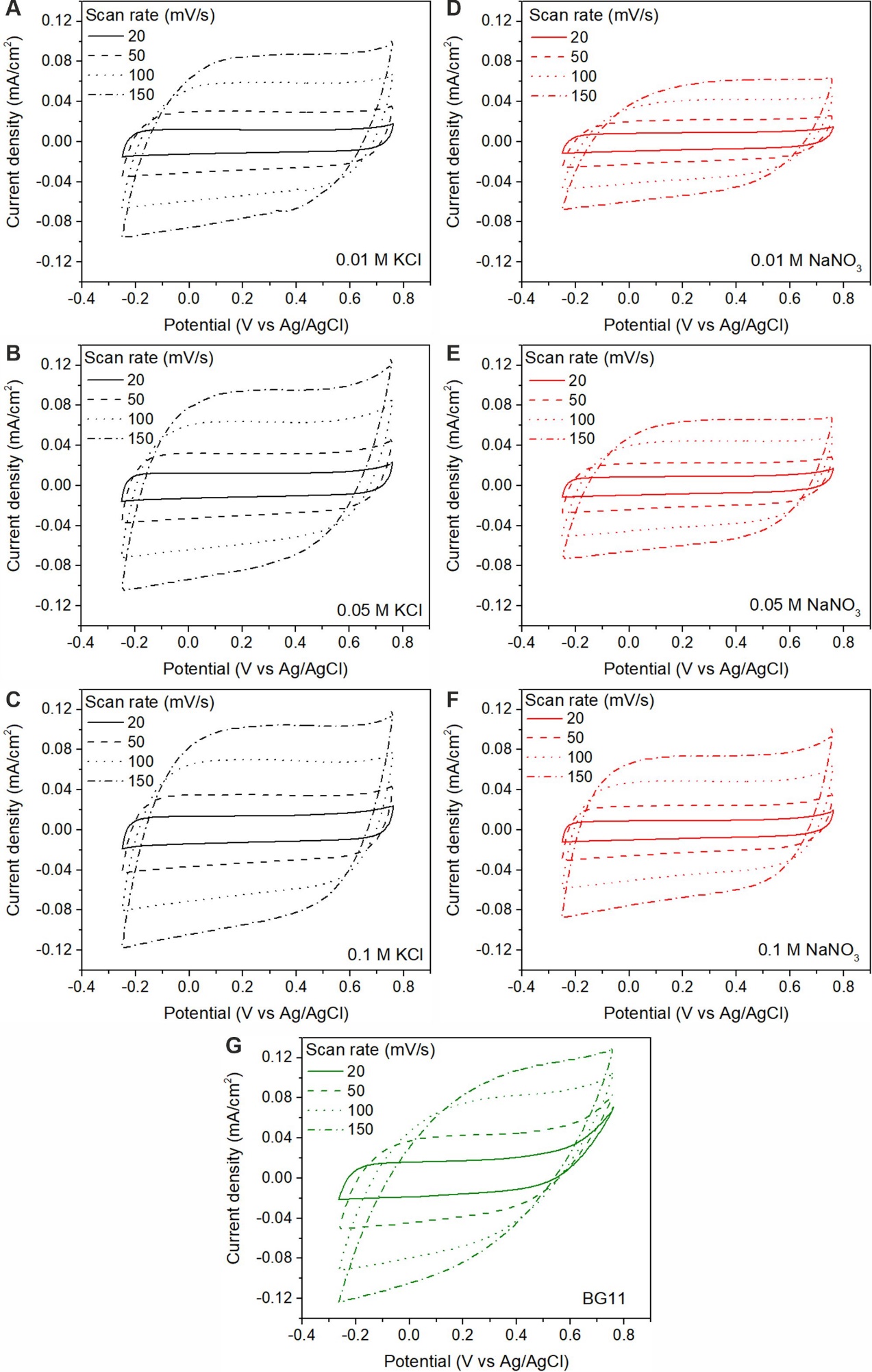

**Supplementary Figure** **2** Cyclic voltammograms of AP-VPP-fabricated PEDOT electrode in different electrolytes. **A)** 0.01 M KCl, **B)** 0.05 M KCl, **C)** 0.1 M KCl, **D)** 0.01 M NaNO_3_, **E)** 0.05 M NaNO_3_, **F)** 0.1 M NaNO_3_, and **G)** BG11 at the scan rate of 20, 50, 100, and 150 mV/s.

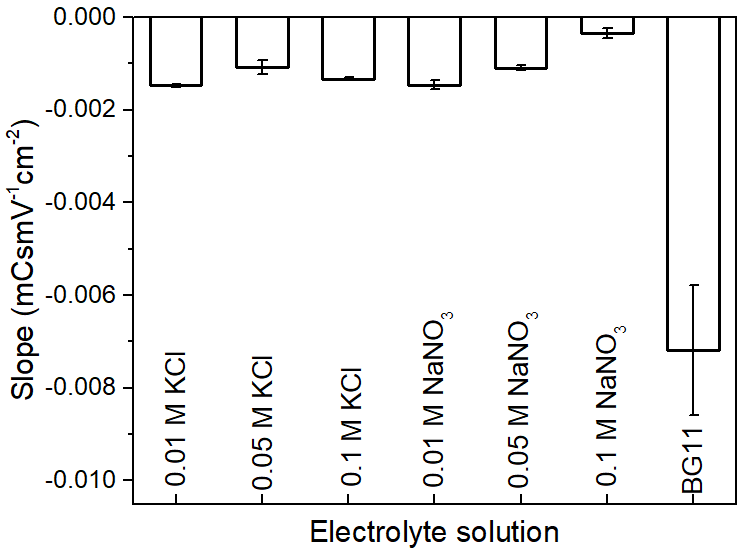

**Supplementary Figure 3** The bar chart shows the tendencies in slopes of linear fitted lines (in Figure 2B, of charge densities against scan rate plots) for each electrolyte and its concentration. The Y error bars show the standard error of the slope of a linear regression.

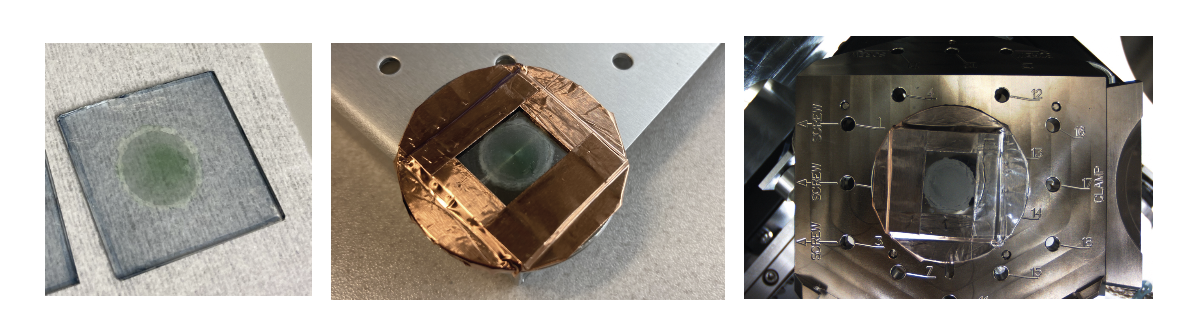

**Supplementary Figure 4** Preparing the bio-anodes for scanning electron microscopy. Left: the biofilm on the electrode after the washing and drying steps. Middle: mounted with copper tape. Right: spray coated with platinum.

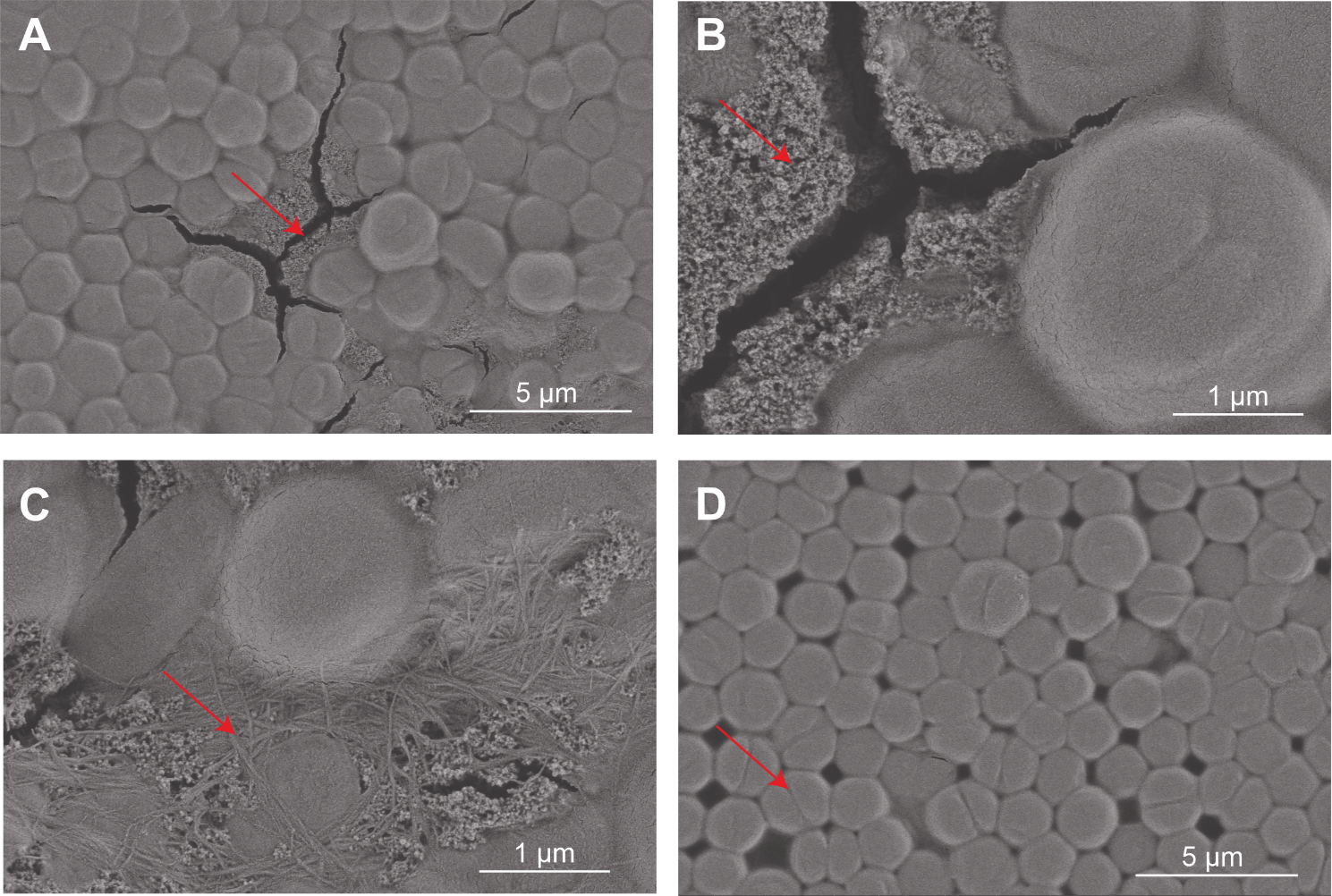

**Supplementary Figure 5** Additional scanning electron microscope images of the bio anode showing features.
**A**) Some cracks in the biofilm (red arrow) typically observed where cells have been displaced. **B)** Extracellular matrix of exopolysaccharides coating the PEDOT electrode surface (Red arrow). **C)** Type IV pili (red arrow). **D)** Dividing cells (red arrow).

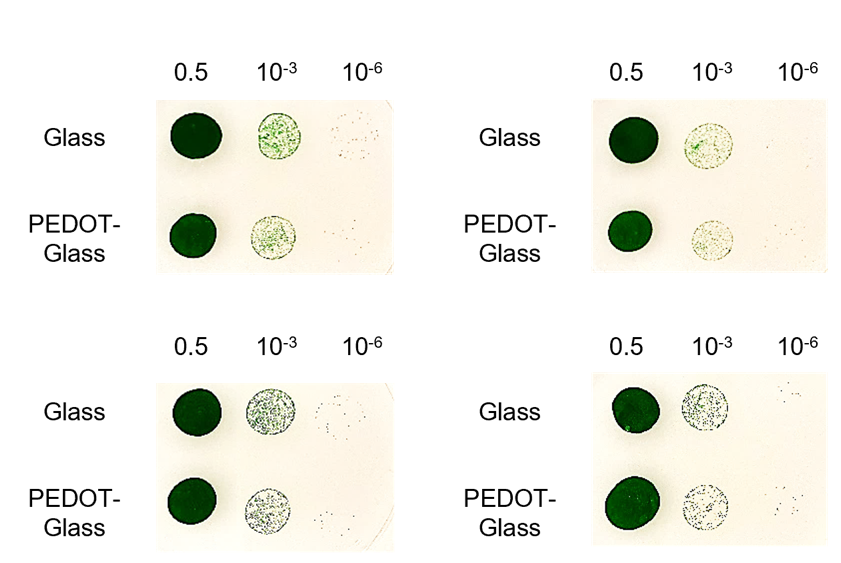

**Supplementary Figure 6** Biocompatibility of AP-VPP PEDOT electrodes with *Synechocystis* sp. PCC 6803 cyanobacterial cells. Column headings are OD_750_ of cell mixture and serial dilutions. Photographs of cells grown from exposure on electrodes. Four biological replicates.

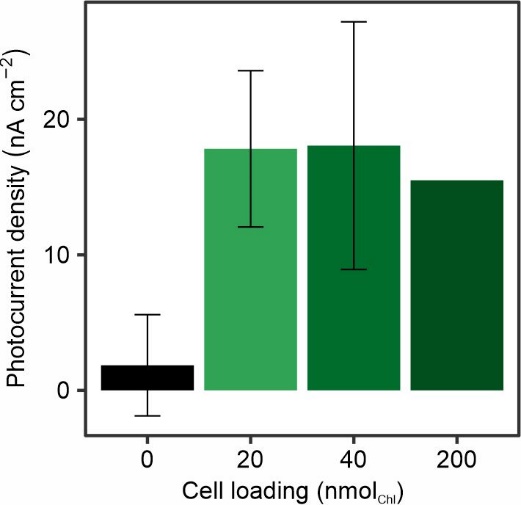

**Supplementary Figure *7*** Effect of different cell loading on photocurrent outputs of Synechocystis cells on AP-VPP-fabricated PEDOT electrodes. Photoelectrochemical parameters: single layer AP-VPP-fabricated PEDOT electrodes, chronoamperometry at 0.1 V vs Ag/AgCl applied potential, 150 µmol photons m^−2^ s^−1^ 460 nm light, 2 min/2 min light/dark cycles. Data presented as the mean ± standard error of the mean of n = 3 biological replicates for 0 and 40 nmol_Chl_ loading, n = 9 for 20 nmol_Chl_ loading and n = 1 for 200 nmol_Chl_ loading.

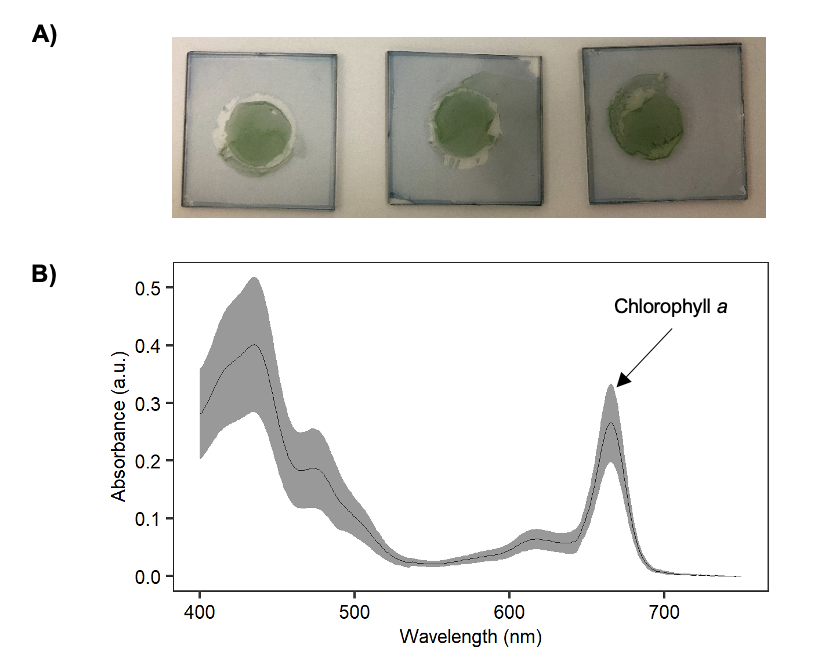

**Supplementary Figure 8** Cyanobacterial cell adherence on AP-VPP-fabricated PEDOT electrodes. **A)** Photographs of biofilms of *Synechocystis* cells on AP-VPP-fabricated PEDOT electrodes after photoelectrochemistry experiments. 20 nmol_Chl_ cells was loaded, 1 layer PEDOT, BG11 (pH 7.5) electrolyte, no mediators or inhibitors were added. The clear glass ring was from the O-ring in the device upon disassembly. **B)** UV-Vis absorbance spectra of the biofilms resuspended in methanol. Data presented as the mean, shading is the standard deviation of three replicates. The chlorophyll a (Chl a) amount was determined using the absorption coefficient of Chl a in methanol at 665 nm (arrow), which is 79.95 Chl a mg ml^-1^ cm^-1^. The Chl a loading was calculated to be 1.86 ± 0.39 nmol.

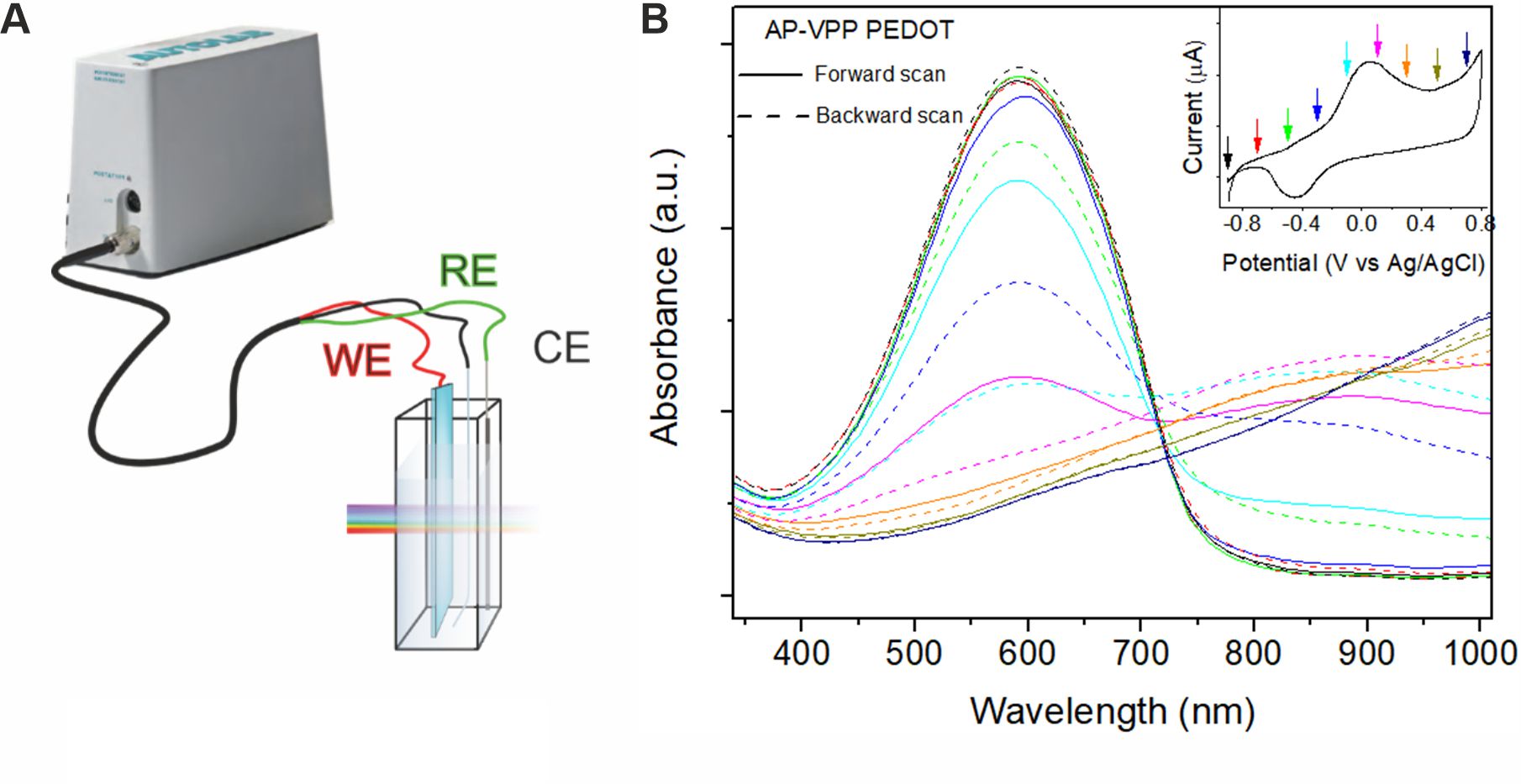

**Supplementary Figure 9** **A)** Schematic of *in situ* UV-Vis spectroelectrochemistry experimental setup (WE: working electrode, RE: reference electrode, and CE: counter electrode) **B)** *In situ* UV-VIS spectra of an AP-VPP-fabricated PEDOT electrode, insert showing CV recorded during in situ UV-VIS measurements (CV measurements performed in 0.1 M tetrabutylammonium tetrafluoroborate (TBA-BF_4_) in acetonitrile, in the range - 0.9 V to 0.8 V at 2 mV s^-1^ scan rate).

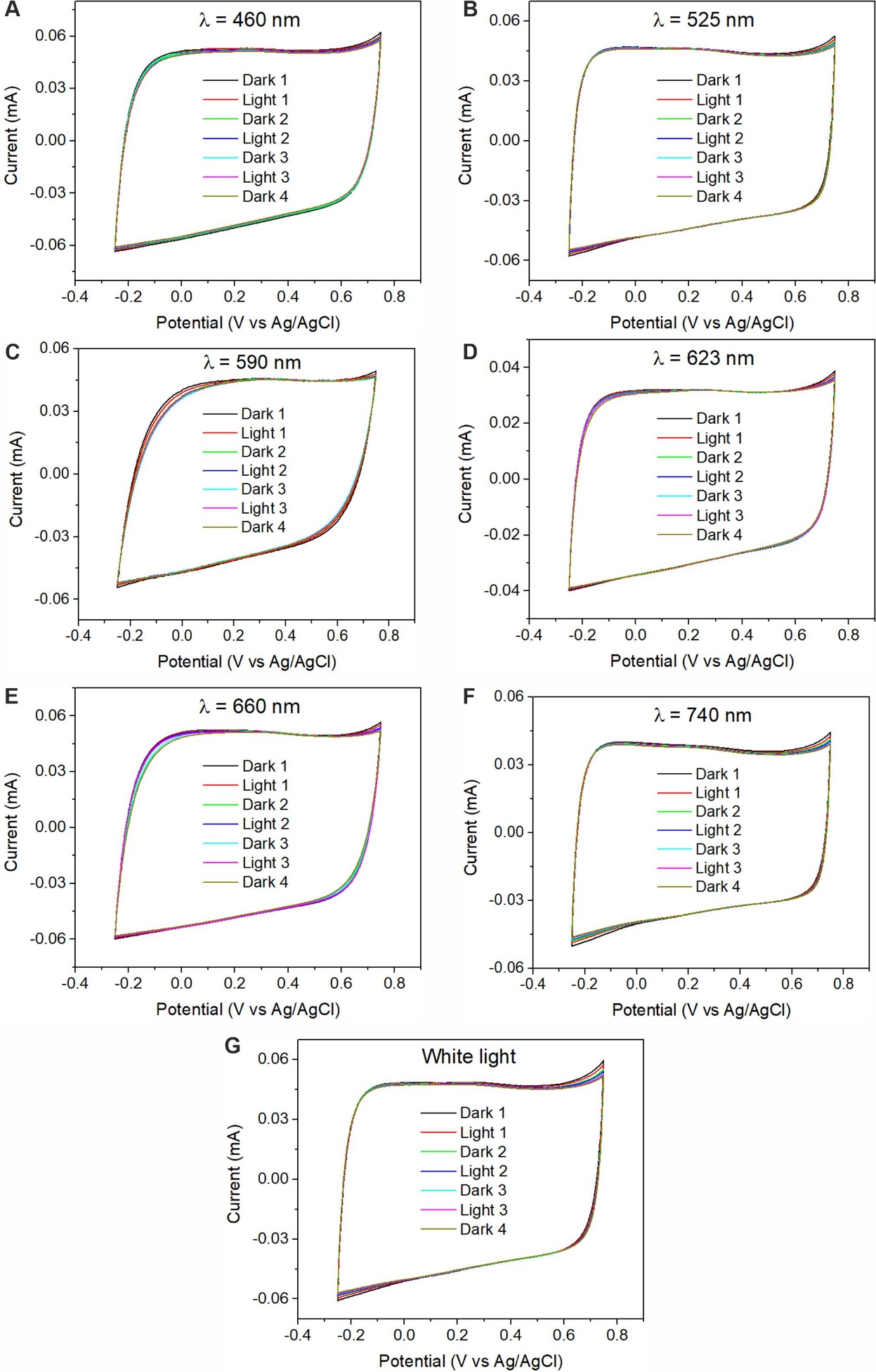

**Supplementary Figure 10** Electrochemical performance of AP-VPP-fabricated PEDOT electrodes under illumination of light at different wavelengths. Photoelectrochemical parameters: single layer AP-VPP-fabricated PEDOT electrodes, no cells loaded, 0.1 M KCl electrolyte, scan rate 100 mV s^-1^, second CV scan shown from dark-light cycles. Light intensities maximum for each wavelength (Supplementary Table 3). Each panel is a different film.
